# Classifying interactions in a synthetic bacterial community is hindered by inhibitory growth medium

**DOI:** 10.1101/2022.03.02.482509

**Authors:** Andrea R. Dos Santos, Rita Di Martino, Samuele Testa, Sara Mitri

## Abstract

Predicting the fate of a microbial community and its member species relies on understanding the nature of their interactions. However, designing simple assays that distinguish between interaction types can be challenging. Here, we performed spent media assays based on the predictions of a mathematical model to decipher the interactions between four bacterial species: *Agrobacterium tumefaciens* (*At*), *Comamonas testosteroni* (*Ct*), *Microbacterium saperdae* (*Ms*) and *Ochrobactrum anthropi* (*Oa*). While most experimental results matched model predictions, the behavior of *Ct* did not: its lag phase was reduced in the pure spent media of *At* and *Ms*, but prolonged again when we replenished with our growth medium. Further experiments showed that the growth medium actually delayed the growth of *Ct*, leading us to suspect that *At* and *Ms* could alleviate this inhibitory effect. There was, however, no evidence supporting such “cross-detoxification” and instead, we identified metabolites secreted by *At* and *Ms* that were then consumed or “crossfed” by *Ct*, shortening its lag phase. Our results highlight that even simple, defined growth media can have inhibitory effects on some species and that such negative effects need to be included in our models. Based on this, we present new guidelines to correctly distinguish between different interaction types, such as cross-detoxification and cross-feeding.

## Introduction

As they grow, microbes modify their environment. This affects other organisms living in their proximity, resulting in measurable interactions. How to classify microbial interactions has been a subject of some debate, but broadly they can be cooperative, competitive or neutral based on one species’ positive, negative or absent effects on another species’ growth, respectively (1).

The way in which such positive and negative effects are physically and chemically mediated may affect the survival of the interacting species (their ecology) and how selection acts on their traits (their evolution) (1–4). Positive interactions occur when one species improves the environment for another, either by reducing its adverse effects or by producing compounds that enhance the other’s growth (1). Whether these improvements also benefit the acting species and/or are costly can affect evolutionary dynamics. For example, siderophores or nutrient-degrading enzymes are useful to their producers as well as other species, but are quite costly (5). Non-producing mutants can then invade the population of producers and destabilize the interaction. But positive interactions can remain stable over time if they are not exploitable: a species may take up nutrients that alter the pH to another species’ benefit (6, 7), or secrete cost-less metabolic byproducts, which can be cross-fed by other co-inhabiting species (8, 9).

Predicting the long-term fate of competitive interactions is equally mechanism-dependent. Competition can be due to one species enhancing harmful conditions (e.g. production of bacteriocins), or removing beneficial ones (e.g. competition for nutrients) (1, 10). Under the latter, known as “exploitative competition”, species compete for resources, and we expect them to evolve to occupy separate niches and compete less (11–14). Under more aggressive “interference competition”, the production of toxins, antibiotics or phage-like particles may result in arms races and species extinctions (10, 15). Competitive interactions often rely on direct cell-to-cell contact (15). In sum, even if positive and competitive interactions are easily measurable at the population level, understanding the mechanisms underlying these measured effects can change the predictions of long-term dynamics or environmental changes.

In natural communities, interactions occur simultaneously between many species, with little evidence of which molecule was produced or consumed by which species, and which species it affects in which way. Identifying interactions and their molecular mechanisms in such complex webs is clearly quite challenging, but can possibly be achieved in a high-throughput manner using spent media (SM) assays, which we will show can distinguish between interaction types without needing to distinguish interacting species, e.g. by fluorescently labelling them. To develop and test the utility of these assays, small synthetic microbial ecosystems of up to a few dozen species are more practical (16–20): Inter-species interactions are easier to disentangle and control, especially since the chemistry of the environment can be designed, and community members can be genetically engineered or selected to exhibit specific interactions (21–26). Their simplicity also allows for parameter estimation in mathematical models to predict community dynamics (22, 27–30).

Here, we aimed to decipher the interactions in a synthetic community we studied previously, composed of four bacterial species: *Agrobacterium tumefaciens* (*At*), *Comamonas testosteroni* (*Ct*), *Microbacterium saperdae* (*Ms*), and *Ochrobactrum anthropi* (*Oa*) (31). This community was dominated by positive interactions when grown in a chemically complex environment. Here, we sought to provide a more controlled environment and used a defined minimal medium (MM) to study the mechanisms behind the interactions between the four species. As we were initially interested in chemical and metabolic interactions that do not require cell-to-cell contact, we grew each species in the SM of all others using an experimental design that allows us to distinguish between interaction types (e.g. exploitative vs. interference competition). Ideally, these simple assays would suffice to identify all types of pairwise interactions, without the need for detailed chemical analyses of secreted and consumed molecules.

We found two strong positive interactions mirroring our previous work: the pure SM of *At* and *Ms* shortened the lag phase of *Ct*. However, we were surprised that this positive effect was lost and even reversed if the spent media were replenished with the original growth medium (MM). Further investigation revealed that the no-carbon (NC) compounds (that cannot be used as carbon sources) in the replenished SM delay *Ct*’s growth despite the presence of enough available carbon sources and the SM. We then wondered whether *At* and *Ms* might remove the inhibitory compounds from the environment for *Ct*, allowing it to grow sooner. Although such cross-detoxification seemed like the most parsimonious explanation, we found no evidence to support it. Instead, using untargeted liquid chromatography–mass spectrometry (LC-MS), we identified at least three molecules secreted by *At* and/or *Ms* that could be metabolised by *Ct* and shortened its lag phase. Our findings show that pinpointing the nature of positive interactions can be quite challenging, as different effects can be intimately intertwined: cross-feeding could alleviate the negative effects of a challenging environment.

## Results

### *At* and *Oa* responded to others following anticipated scenarios

To identify the interactions between the four species in our simplified medium, we conducted spent media (SM) assays by growing each species alone until stationary phase in a minimal medium (MM) containing a no-carbon part (NC = salts and trace metals) and two carbon sources (CS): glucose and citric acid (MM = NC + CS, see Methods). We then removed the cells by filtration and grew each species in all other resulting spent media (SM). We compared the growth in SM (condition IV) to the growth in a medium comprising only the NC compounds (condition I), a positive control of MM (condition II) and two other conditions (Fig. 1A, left): First (condition III), we diluted (1:1) the SM in 2× NC so that the species have access to at least the same concentrations of salts and trace metals as in the fresh MM, and second (condition V), SM was diluted (1:1) in 2× MM to contain at least the original concentration of carbon sources, salts and trace metals.

**Fig. 1.**
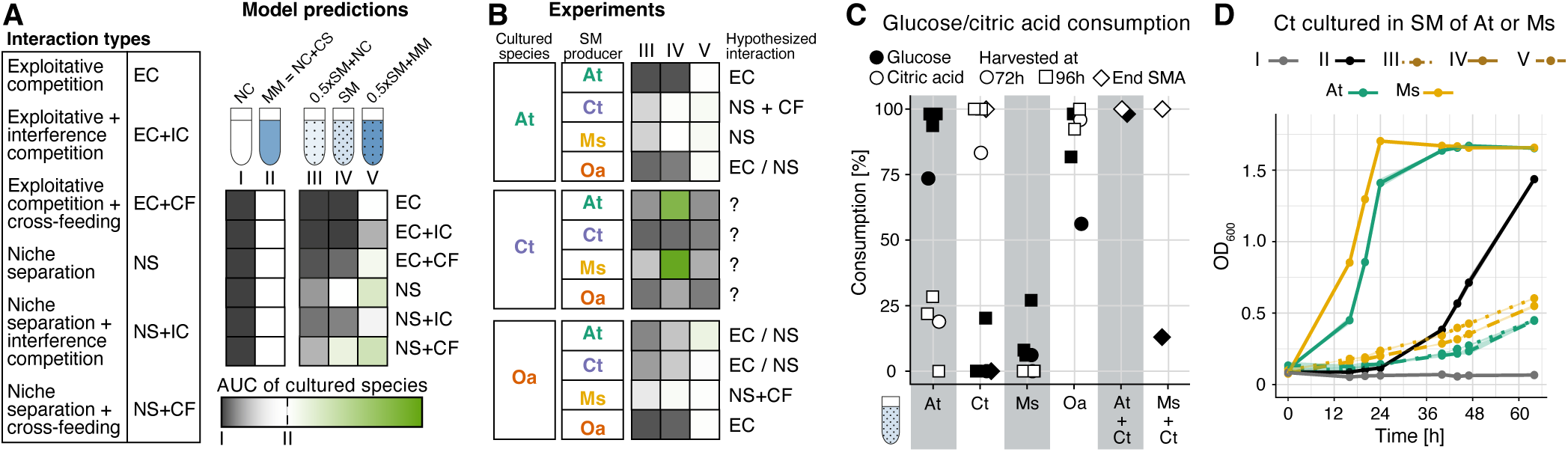
Spent media assays. (A) We used our model to simulate the growth of species under different types of positive or negative interactions, as listed in the table and illustrated in Fig. S1. Five simulated growth conditions allowed us to distinguish between these four different interaction types: I. no-carbon (NC) medium containing only salts and trace metals, but no carbon sources, II. medium I but with 15 mM glucose and 10 mM citric acid (CS: carbon sources), III. a mix of the spent medium of a given partner species (50%) and medium I (50% of a 2× concentrated solution), IV. the spent medium of a partner species, V. a mix of the spent medium of a partner species (50%) and medium II (50% of a 2× concentrated solution). Square colors reflect area under the growth curve (AUC) under these different conditions relative to the AUC in conditions I and II (gradient legend). See Fig. S2 for model growth curves that generated these. (B) Results of spent media (SM) assays, where each row shows how a given species grew normalized by conditions I and II in the different SM conditions (III, IV, V) for each SM producer (see Fig. S3 for growth curves and Fig. S4 for AUCs with statistical comparisons). Comparing these patterns to the predictions in panel A determined the hypothesized interactions. When *Ct* was grown in the spent media of others, interactions did not qualitatively match any predicted scenarios. The growth of *Ms* is not shown here, as the OD data and the CFUs data were not coherent. (C) Consumption [%] of glucose and citric acid by the 4 species based on commercial chemical kits applied to the spent media in stationary phase (72 and 96 hours of growth). We also analyzed the SM of *Ct* after growing in the pure SM of *At* and *Ms* (samples collected at the end of the SM assay after ∼ 64 hours, thus labelled “end SMA”). (D) Growth curves of *Ct* as OD_600_ over time when grown in the five culture conditions, with either *At* or *Ms* generating the spent media, (n = 3, *±*sd, error bars are present but very small). Roman numerals indicate culture conditions as in panel A, colours indicate spent media producing species.

Our expectations for these five different experimental conditions were established using a mathematical model where we simulated the outcome of six interaction types: exploitative competition, exploitative with interference competition, exploitative competition with cross-feeding, niche separation, niche separation with interference competition, and niche separation with cross-feeding (Figs. S1, S2). We calculated the area under each simulated growth curve, and compared it to the negative and positive controls (I and II) (Fig. 1A). For example, if two species compete for the same carbon source or a species is grown in its own SM (exploitative competition, EC), the carbon source in the model is depleted such that condition III and IV are similar to I. The replenishment of the carbon sources in condition V restores growth to the level of the positive control II. The remaining base expectations are shown in Figs. 1A and S2.

The spent media experiments were analyzed by calculating the area under the OD_600_ growth curves. *At* and *Oa*’s growth could be classified according to the six anticipated scenarios (Fig. 1B, S3, S4). We have omitted data on *Ms*, as quantifying its growth was problematic due to contradictions between our measurements (Fig. S3, S5). Surprisingly, the growth patterns of *Ct* did not correspond to any of the expected scenarios (Fig. 1B).

We first verified the behavior of *At* and *Oa* by quantifying which carbon sources the four species use (see Methods). We found that *At* consumes mostly glucose, *Ct* mostly citric acid, *Oa* consumes both, while *Ms* consumes little of either (Fig. 1C). In agreement with this, *At* reflects the niche separation model in the spent media of *Ct* and *Ms*. Since *At* does not consume citric acid, but overlaps with *Oa* in consuming glucose, SM interactions of both *At* and *Oa* lie in between exploitative competition and niche separation. Similarly, in the SM of *Ct, Oa* follows a pattern in between exploitative competition and niche separation. We observed some evidence for cross-feeding from *Ms* to *Oa*, and for interference competition from *Ct* to *Oa* (see Fig. S4 for statistics), but do not explore this further. Instead, we focus on why *Ct* did not fit our model’s predictions.

### *Ct* has a shorter lag phase in the pure spent media of *At* and *Ms* but grows poorly in all other conditions

When *Ct* grows in the SM of either *At* or *Ms* (condition IV), its AUC is significantly higher (t-test with Bonferroni correction, both *df* = 5, *P <* 0.001) than in II, which is due to a much shorter lag phase (Fig. 1D). However, when we replenished the growth medium (condition V), *Ct* grew significantly worse than in the original medium (condition II) (both *df* = 5, *P* < 0.001, Fig. 1B, D). We observe the same pattern in its own and in *Oa*’s SM (growth in V is significantly worse than II, *P <* 0.001, Fig. S4). This is surprising because condition V should contain at least the same concentration of carbon sources as in condition II.

This led us to suspect that some of the replenished compounds might impair the growth of *Ct* because they end up at higher concentrations than in the original medium. Accordingly, we tested a first hypothesis: that one or more of the compounds in the NC medium inhibit the growth of *Ct*, while *At* and *Ms* are able to reduce their concentration and thereby shorten the lag phase of *Ct*. According to our model, such “cross-detoxification” could be a valid explanation for the observed patterns (Fig. S6).

### No-carbon compounds delay the growth of *Ct* but are not reduced by *At* or *Ms*

To explore whether the NC medium could affect the growth of *Ct*, we first manipulated its two main ingredients: M9 and Hutner’s vitamin free Mineral Base (HMB, see Methods). While reducing the concentration of HMB had no positive effect (Fig. S7), a small decrease in the concentration of M9 shortened the lag phase of *Ct*, while reducing it further had a detrimental effect (Fig. 2A, see Fig. S8A for CFUs). As changing the concentration of M9 changes both pH and osmolarity simultaneously, we next tested the effect of each separately and found that both (i) increasing the pH and (ii) decreasing M9 but keeping the pH constant (lowering osmolarity) shortened the lag phase of *Ct* (Fig. 2B). To assess if specific compounds in the M9 (see Methods) could influence the lag phase of *Ct*, we tested the effect of each compound on the growth of *Ct* by decreasing their concentration in the NC (Fig. 2C). Of all the compounds, we found that the total concentration of phosphate (Na_2_HPO_4_ and KH_2_PO_4_) was the only one that influenced the lag phase of *Ct* independently of the pH (Fig. 2D). Changing the ratio of the two ions (Na^+^ and K^+^) by changing the ratio of the corresponding salts (Na_2_HPO_4_ and KH_2_PO_4_) also affected the lag phase: a higher proportion of potassium shortened it while sodium increased it (Fig. 2E). These findings suggest that the NC medium is suboptimal for *Ct* and lengthens its lag phase: a controlled pH, a lower phosphate concentration, or a smaller sodium to potassium ratio in the M9 allow *Ct* to grow earlier. Accordingly, our most parsimonious explanation for its SM behavior (Fig. 1B) was that *At* and *Ms* can modify at least one of those factors in the medium. However, when we measured pH, osmolarity, and the concentration of phosphate, and sodium and potassium ions in the SM of the 4 species (Fig. S9), we found that none differed between the original MM and the SM of *At* and *Ms* in a way that would explain the shortened lag phase of *Ct*. In sum, several properties of the NC medium lengthen the lag phase of *Ct*, but *At* and *Ms* appear unable to significantly modify these properties in a way that would explain why *Ct* grows so well in their SM. We therefore rejected our first hypothesis.

**Fig. 2.**
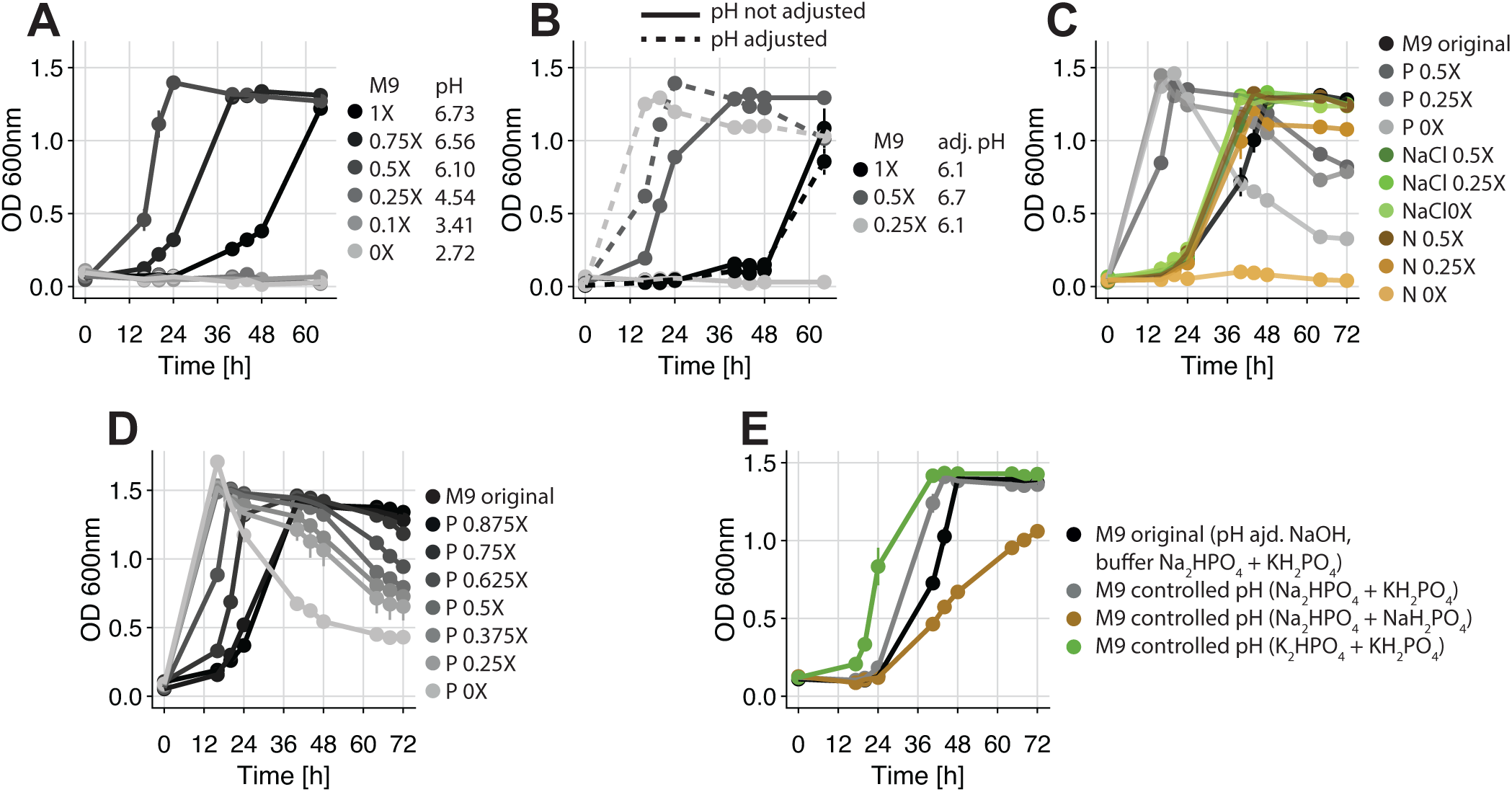
Growth of *Ct* (as OD_600_) in minimal medium (MM) made with no-carbon (NC) medium where we modified the M9 part. (A) Growth in MM with decreasing concentrations of M9. The resulting pH is indicated for each fresh MM, showing that decreasing the concentration of M9 also decreases the pH. (B) Growth in MM with decreasing concentrations of M9 with and without manual pH adjustment to either 6.1 or 6.7 using NaOH. Decreasing the concentration of M9 while keeping a pH close to neutrality shortens the lag phase of *Ct*. (C) Growth in MM with M9 that vary in the concentration of its main components: phosphate (overall Na_2_HPO_4_ plus KH_2_PO_4_ concentration, abbreviated P), sodium chloride (NaCl) and nitrogen (NH_4_ Cl, abbreviated N). The pH is adjusted to 6.7 with NaOH in all conditions (pH of the original MM). The overall concentration of phosphate seems to be the sole factor that affects *Ct* ‘s lag phase (D) Growth in MM with M9 varying in its overall concentration of phosphate (Na_2_HPO_4_ plus KH_2_PO_4_, abbreviated P). The pH is adjusted to 6.7 with NaOH in all conditions (pH of the original MM). (E) Growth in MM with M9 that have either phosphate compounds comprising only Na^+^ ions (Na_2_HPO_4_ plus NaH_2_PO_4_), only K^+^ ions (K_2_HPO_4_ plus KH_2_PO_4_) or both ions (Na_2_HPO_4_ plus KH_2_PO_4_) but with adjusted ratios so that the pH is at 6.7 (“controlled pH”) with no NaOH adjustment. For all graphs: mean is plotted *±*sd, n = 3.

### *Ct* feeds on metabolites produced by *At* and *Ms*

Another hypothesis that could explain the shortened lag phase of *Ct* in the SM of *At* and *Ms* is that their SM contains metabolic by-products that allow *Ct* to grow earlier. To find such candidate molecules, we performed an untargeted Liquid Chromatography - Mass Spectrometry (LC-MS) analysis (see Methods) on the SM of *At, Ct* and *Ms* and on the SM of *At* and *Ms* after *Ct* had grown in it to assess if *At* and *Ms* secrete molecules that *Ct* then consumes (Fig. 3A, full list in Fig. S10).

**Fig. 3.**
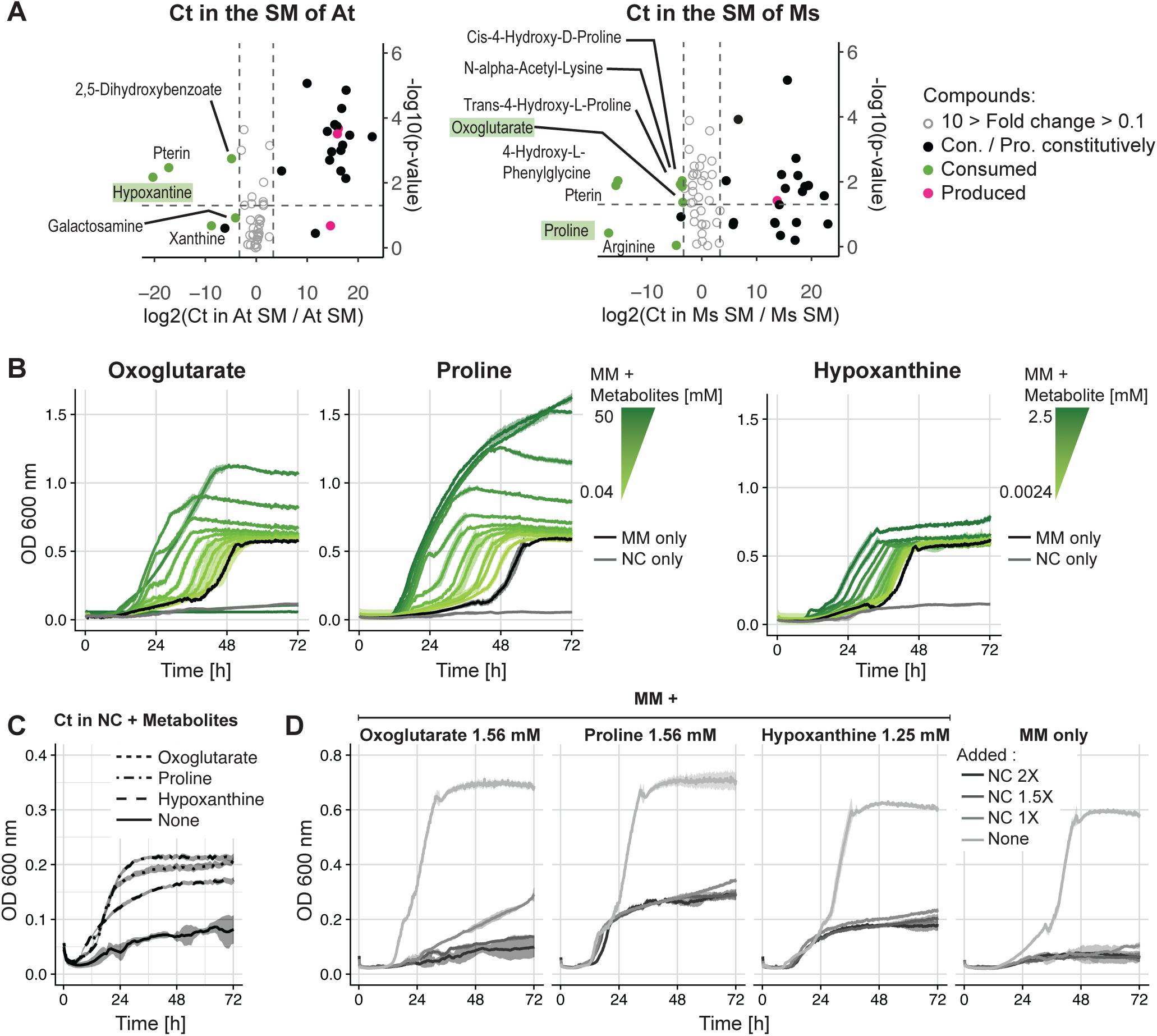
Identification of three cross-fed metabolites (oxoglutarate, proline, hypoxanthine) and their effect on *Ct*. (A) Untargeted metabolomics analysis (focusing on polar molecules) performed on the SM of *At, Ct* and *Ms* and on the SM of *At* and *Ms* after *Ct* grew in them to identify metabolites that were produced by *At* and *Ms* to then be consumed by *Ct* (relative quantification). Left: X-axis shows the log_2_ of the ratio (fold change) between the abundance of each metabolite in the spent media of *At* after the growth of *Ct* to the abundance of each metabolites in the spent media of *At*. If metabolites are on the left of the graph (negative), *Ct* consumed them from At’s SM, while if they are on the right (positive) *Ct* produced them. Grey metabolites between the dashed lines have a fold change that is *<* 10×, which we did not consider. Y-axis shows the log_10_ of the p-value, the 0.05 significance threshold is represented by a dashed line. Metabolites in green boxes were chosen for further analysis. Right: Same as the left panel but for the SM of *Ms*. In both panels, n = 3. ANOVA one-factor (on log10 transformed data) was used to test the significance of metabolite changes in the different conditions. (B) Effect of oxoglutarate, proline and hypoxanthine on the growth of *Ct* in a range of concentrations added to MM, measured as the OD_600_ over 72 hours. The mean is plotted and the transparent area around the curves represents the standard deviations. As a control, *Ct* was grown in MM or NC, (n = 4). (C) Effect of 1.56 mM oxoglutarate, 1.56 mM proline and 1.25 mM hypoxanthine on the growth of Ct in NC medium only, with NC alone as a control over 72 hours. The OD was measured every 10 minutes, the mean is plotted and the transparent area around the curves represents the standard deviations, (n = 3).The scale of the y-axis is smaller that in panel D to better show the growth curves. (D) Effect of intermediate concentrations of each metabolite in MM with increasing concentrations of NC (50% replenishment in either NC medium 1×, 1.5×, or 2×), measured as the OD_600_ over 72 hours. The mean is plotted and the transparent area around the curves represents the standard deviations. As a control, *Ct* was grown in MM (in the same conditions, (n = 3). The data show that all three metabolites could act as carbon sources, could short *Ct* ‘s lag phase and the effect was reversed on the addition of NC.

We identified several compounds that follow this pattern (Fig. 3A) and selected three – based on availability, cost and ease of use – to assess their effect on the lag phase of *Ct*: hypoxanthine, oxoglutarate and proline (Fig. 3B). As the LC-MS only yielded the relative abundances of each compound, we added several concentrations to *Ct* growing in MM. We found that a range of concentrations shortens the lag phase of *Ct* significantly (Oxoglutarate: from 0.39 mM to 12.5mM, table S1; proline: *≥*0.39mM, table S2; hypoxanthine: *≥*0.62mM, table S3). At high enough concentrations, all three metabolites also increased the final yield of *Ct*, suggesting that they act as carbon sources (Oxoglutarate: from 0.09 mM to 12.5 mM, table S4; proline: *≥* 0.78 mM, table S5; hypoxanthine: *≥* 1.25 mM, table S6). At its two highest concentrations (25 and 50mM), oxoglutarate even had an inhibitory effect on *Ct*. To confirm that these three metabolites could act as carbon sources, we cultured *Ct* in NC medium containing each of the metabolites at an intermediate concentration (oxoglutarate and proline: 1.56mM; hypoxanthine: 1.25mM) as the sole carbon source and observed significant growth in all three cases (Fig. 3C).

It appears then that these three metabolites are being crossfed by *Ct*. If they are responsible for the shortening of the lag phase of *Ct* in the SM of *At* and *Ms*, the replenishment of the medium should cancel this effect like we observed in the spent media assays (Condition III, V, Fig. 1B, D). We grew *Ct* in MM supplemented with each of the three metabolites (at concentrations based on Fig. 3B) and added increasing concentrations of the NC medium (Fig. 3D). As in our original experiments, the growth of *Ct* was impaired despite the presence of the metabolites and the original carbon sources in all conditions compared to the positive controls (no addition of NC or metabolites), supporting our hypothesis. These results are in line with our updated model (Fig. S6) that includes the inhibitory effect of the medium.

Overall, we find that different properties of the NC compounds can lengthen the time *Ct* takes to start growing. Metabolites secreted by *At* and *Ms* – at least the three that we tested – are able to reduce this effect through cross-feeding. However, when the concentration of NC compounds is high enough, the effect of the metabolites is insufficient and *Ct* grows very little on the timescale of our experiments.

## Discussion

Before running our SM assays, we used a simple mathematical model to generate our base expectations under different experimental conditions. We were surprised when the behavior of one species, *Ct*, did not correspond to the predictions of any of the scenarios simulated by the model. Our further analysis revealed that this was because our model assumed that the growth medium could either allow cells to grow or not, but it could not inhibit species, such that increasing its concentration would delay their growth. In hindsight, we know that designing growth media for different species can be quite challenging (32, 33), so we might expect negative effects of some media on some bacterial species.

Based on this new intuition, we updated our model to cover cases when the medium inhibits growth (Fig. 4A, Fig. S6). The patterns in this new model fit qualitatively with what we observed for *Ct*, but also drew our attention to another issue: it was challenging to distinguish between cross-feeding and cross-detoxification. And even though cross-detoxification seemed the more parsimonious explanation, all our experiments led us to reject it as the underlying interaction. The new model does, however, give two clues to distinguish these two types of positive interaction without chemical analyses: First, in the pure spent media (condition IV), the final yield in a cross-detoxification interaction will never exceed that of the original growth medium (under the assumption that inhibition slows down growth rather than increasing death), while this should be the case for cross-feeding (right arrow in Fig. 4B, C, S6); second, in cross-feeding, one may detect a first short stationary phase as cells switch their metabolism to consume the second carbon source (left arrow in 4B). We reran our original experiment in a plate reader to obtain higher time resolution, and indeed found a significant difference in the final yield of *Ct* in MM and the SM (condition IV) of *At* (*P <* 0.01) and *Ms* (*P <* 0.001), as well as a small “bump” at the beginning of the SM growth curves (arrows in Fig. 4D, Fig. S11, see tables S7 and S8 for statistics on the final yield and the length of the lag phase, respectively).

**Fig. 4.**
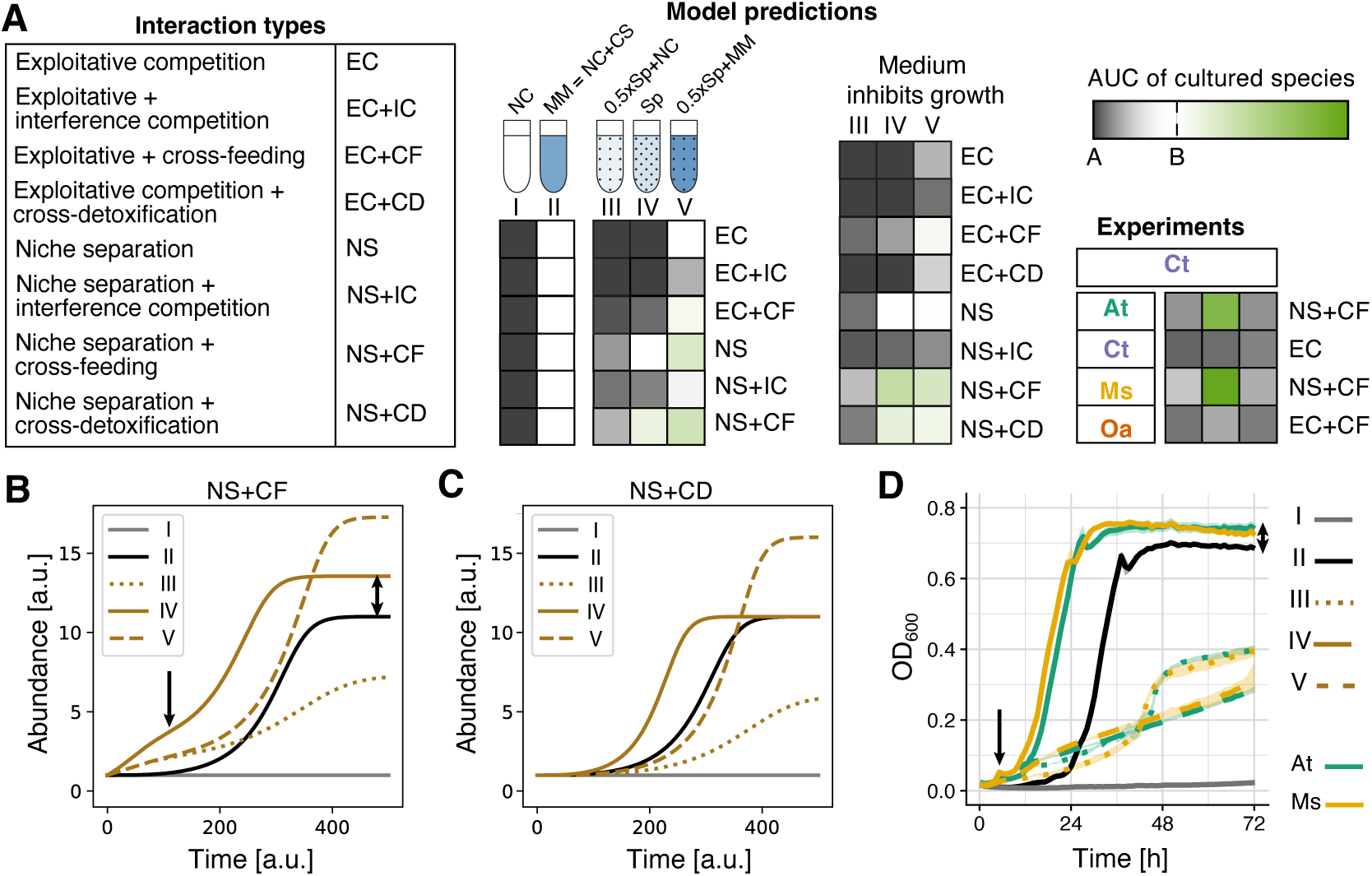
Revised model better predicts *Ct* ‘s growth patterns. (A) Model predictions of the original model on the left (same as in Fig. 1A), the model predictions in an environment that inhibits growth in the center and the data for the growth of *Ct* in the SM of the four different species. Although the predictions do not match perfectly, they qualitatively resemble the patterns in the inhibitory growth medium. (B) Model predictions for niche separation + cross-feeding. The single arrow shows the growth “bump” in IV compared to II at around 80 hours due to a switch in carbon sources, and the double arrow a greater final yield in condition IV. (C) Model predictions for niche separation + cross-detoxification. Here, the bump is missing and the final yield is identical in II and IV. (D) Repetition of the experiment shown in Fig. 1D in a 96-well plate with higher time resolution shows the presence of a bump in the two species’ SM (condition IV, solid yellow and green lines), indicating a switch in carbon source (see Fig. S11), and a greater final yield compared to growth in fresh medium (condition II, black). Note that the model still does not correspond perfectly to the experimental data, indicating that there may be additional interactions at play.

Our simple spent media assays and updated model can in principle suffice to classify the dominant interactions, at least in defined medium with few carbon sources. They are more powerful than direct co-culture experiments, where the directionality of interactions is obscured. In Fig. S12, we have constructed a decision tree that allows one to classify data using the 5 spent media conditions outlined here, assuming that each species consumes a single carbon source, regardless of how many are present. This is because for simplicity, the medium in our model only includes a single carbon source (it can be extended). We therefore do not apply the decision tree to interactions with *Oa*, which we classified based on our knowledge that it consumes both carbon sources.

Even in the simple scenario we have studied, we may be missing additional layers of interactions between these four species. First, all three SM metabolites we tested turned out to be cross-fed, suggesting that others may have a similar effect. This aligns with previous research suggesting that crossfeeding might be quite common (34, 35), particularly for species that are in a sub-optimal environment that contains few carbon sources (36, 37), or that is toxic (31). It may be then, that the difficulty of finding a growth medium that can sustain the individual growth of all members of a community, is precisely what allows us to observe cross-feeding interactions. In fact, positive interactions are often “accidental”, resulting from the secretion of cost-less metabolic by-products by the few species that are able to grow (8, 19, 36, 38–44). While the secretion of metabolites like amino acids by bacteria might seem counter-intuitive, several mechanisms like the maintenance of cell homeostasis (release of over-produced metabolites) or cell lysis can explain costly metabolic byproduct secretion (45).

We were also surprised that *Ms* produced several metabolites (Fig. 3A) and affected *Ct*’s growth, even though its own population size did not increase significantly (Fig. S5). This aligns with other studies showing that cross-feeding does not require bacterial growth (8). Indeed, the absence of growth does not necessarily indicate metabolic inactivity, as metabolic activity is required to produce enough energy for survival. This suggests that in larger bacterial communities, such as the gut microbiome, non- or slow-growing species should not be ignored, as they may still significantly affect other community members.

Despite our efforts, it still remains unclear why the minimal medium delays *Ct*’s growth, and why the cross-fed compounds allowed it to start growing sooner. One hypothesis is that *Ct* experiences osmotic stress in the minimal medium, which can be reflected in the length of its lag phase (46, 47). Given that the other 3 species seemed robust to this stress, the metabolites they secrete could, once consumed, help *Ct* to cope with this stress. Proline, for example, can act as a “compatible solute” (48–51), which are molecules that bacteria synthesize or take up to balance osmotic pressure in hyper-osmotic environments. Alternatively, the metabolites could act as metabolic precursors, allowing *Ct* to synthesize its own compatible solutes *de novo*. This may be the case for oxoglutarate, which is a direct intermediate in the Krebs cycle (52). In *E. coli*, oxoglutarate is taken up from the environment (53), but is also leaked (54), hinting that it may be involved in extracellular exchange. Similarly, the purine derivative hypoxanthine is an important nitrogen source and participates in nucleic acid synthesis via the pentose phosphate salvage pathway (55). Interestingly, hypoxanthine was found to mediate interactions influencing biofilm formation between *B. subtilis* and soil bacteria whose cell-free supernatants were analysed similarly to our own approach (HPLC, NMR and HR-MS) (56). Another hypothesis is that *Ct* requires a metabolic shift to grow on citric acid compared to other carbon sources, and that increases its lag phase due to a high enzymatic cost (57, 58). The presence of metabolites secreted by *At* and *Ms* could then allow it to metabolize citric acid more rapidly. Distinguishing between these different hypotheses could be achieved by engineering *Ct* to report on osmotic stress, by isotopically labelling the carbon sources and following their metabolic by-products from *At* and *Ms* that are later consumed by *Ct*, and/or by testing the role of the remaining identified metabolites.

The ability to classify interspecies interactions to the level of distinguishing cross-feeding from cross-detoxification, for example, is not just a matter of curiosity, but is key to understanding and predicting community dynamics (2, 59, 60). Even mechanistic details of cross-feeding can affect community dynamics. La Sarre et al. (59) showed that increasing the concentration of a cross-fed metabolite can render it toxic to the partner species, leading to a new community equilibrium. To make matters even more complicated, each species pair is likely to interact in more than one way, but the effects we observe are cumulative (60). Here, we showed how *Ms*, for example, produced a whole series of compounds, and that *Ct* could feed on the two that we tested. But it may well be that other compounds have small inhibitory effects on *Ct*, and that changing the environmental conditions could increase their production or leakage rates and alter community dynamics (59).

In conclusion, our work proposes that carefully designed spent media assays together with a simple mathematical model can help to map out the dominant metabolic interactions in more detail than simply labeling them as positive or negative. We have showcased this using a small, synthetic bacterial community in a defined medium, in which we could verify the read-outs from the growth curves with more detailed analyses. It remains to be seen whether our approach would scale up to more high-throughput approaches in larger communities (as in (36), e.g.) and more complex environments. But ultimately, such simple experimental approaches are needed to predict the dynamics of natural microbial communities.

## Material and methods

### Cell culture preparation

Species were grown in monoculture in TSB (tryptic soy broth) overnight (28°C, shaking 200 rpm) and were then diluted to an OD_600_ of 0.05 in fresh TSB and incubated again for 3 hours in order to reach exponential growth. Each culture was then washed 2 times in PBS (phosphate buffered saline) (centrifugation: 15 min at 4’000 rpm, room temperature) and the final bacterial pellets were resuspended in the adequate medium so that the initial OD_600_ would be 0.1.

### Colony forming units (CFUs) measurement

To measure the CFUs, we sampled 20*μ*L of our cultures and diluted them in 180*μ*L of PBS in 96-well plates and proceeded to 10-fold dilutions down to 10^−7^. The dilutions were plated on TSA (tryptic soy agar) as drops and were then spread into lines.

### Media recipes

For the compositions of the different solutions used, see the following tables: minimal medium (MM) - table S9; no-carbon (NC) medium - table S10; HMB - table S11; Metal 44 - table S12; M9 - table S13; M9 with controlled pH used in Fig. 2E - tables S14, S15, S16.

### Spent media (SM) assays

Spent medium (SM) from each of the four species were obtained by growing them in large volumes (V = 30 mL) in MM until they reached stationary phase (∼72h to 96 hours, decided by OD_600_ determination). We then centrifuged the bacterial culture (20 min, 4’000 rpm, room temperature) and collected the supernatants. We centrifuged the supernatants again to eliminate as many bacterial cells and debris as possible before filtering them using vacuum filters (TPP vacuum filtration “rapid”-Filtermax, membrane: PES, 0.22 *μ*m). From those SM we prepared 3 media conditions to test the effect of the SM on our 4 species: conditions C (SM:NC 2× = 1:1), condition D (pure SM only) and condition E (SM:MM 2× = 1:1). Our control conditions were fresh NC medium (A, negative control) and fresh MM (B, positive control). We grew the 4 species in mono-cultures in those 5 conditions (V = 4 mL) over 60 hours and measured the OD_600_ over time (using Ultrospec 10 cell density meter, Biochrom) and performed CFU counts before the initial incubation, at 24h and 48h. We calculated the area under the OD_600_ growth curves (AUC) and used this value as a proxy for growth (Andri et mult. al. S (2021). DescTools: Tools for Descriptive Statistics. R package version 0.99.44) (R version 4.1.2). We repeated those SM assays in 96-well plates (V = 200*μ*L) to increase the resolution of our data and measured the OD_600_ every 10 minutes for 72 hours using a microplate reader (Biotek synergy H1, 28°C, continuous double-orbital shaking).

### Glucose, citric acid, phosphate, sodium, potassium and osmolarity quantification

To quantify glucose, citric acid, phosphate, sodium and potassium in the SM of *At, Ct* and *Ms* we used different chemical kits. We generated SM for each species as described previously (section “Spent media (SM) assays”, total incubation time ≃89 hours) and followed the specific protocols of each kit to determine which concentration of SM to test given the theoreticalconcentration of the tested compounds in fresh MM. **Glucose kit**: Glucose (HK) Assay Kit (Sigma, product Code GAHK-20). **Citric acid kit**: Citric Acid Assay Kit (Megazyme, product Code: K-CITR). **Phosphate kit**: Phosphate Assay Kit (Colorimetric) (Abcam, product code: ab65622). **Sodium kit**: Sodium Assay Kit (Colorimetric) (Sigma, product code: MAK247). **Potassium kit:** Potassium Assay Kit (Fluorometric) (Abcam, product code: ab252904). Osmolarity was measured in each SM sample using an osmometer (Osmomat 030 by Gonotec).

### Metabolomics analyses of SM samples

Untargeted metabolomics analyses were performed for us at Metabolomics Platform, Faculty of Biology and Medicine, University of Lausanne, on the following samples, focusing on polar (water soluble) compounds : fresh minimal medium (MM); SM of *At, Ct* and *Ms* (generated as described in “Spent media (SM) assays”) and the SM of *At* and *Ms* after *Ct* grew in them (total incubation time ≃60 hours). To summarize the procedure, we first identified which metabolites were produced by *At, Ct* and *Ms* when grown in mono-culture in MM (fresh MM compared to the SM of *At, Ct* and *Ms*), and then compared this list of metabolites to the ones identified in the SM of *At*/*Ms* after *Ct* grew in them (SM of *At* and *Ms* compared to the SM of *At* and *Ms* after *Ct* grew in them). Using those comparisons we could identify 64 compounds that were produced by *At* and *Ms* and then consumed by *Ct* or that were absent in the SM of *At* and *Ms* but were later produced by *Ct*. From that list we only considered the compounds consumed by Ct with a fold change of at least 10. Further details are in Supplementary note S2.

### Testing the effect of metabolites on Ct: oxoglutarate, proline and hypoxanthine

We followed a similar protocol for all the compounds. For *oxoglutarate* and *proline*, the same concentration was tested. In a 96-well plate, we added 180*μ*L of water in wells B-E(1) (4 replicates). We then added 20 *μ*L of a 1M stock solution of either oxoglutarate or proline so that those wells contained 100mM of the metabolite tested. The other wells in lines B to E were filled with 100*μ*L of water. Serial dilutions from wells B-E(1) to B-E(11) were performed by transferring 100*μ*L each time (2× dilutions). This way we obtained 11 concentrations to test on *Ct* (from 50mM to 0.04mM), in 4 replicates. To those wells, we then added 100*μ*L of *Ct* cultures that were in MM 2× concentrated and with an OD of 0.2 (following the method described in “Cell culture preparations”). This way, the final concentration of MM is 1× and the final OD is 0.1 (as usually tested). As controls, we grew *Ct* in MM (in 4 wells we mixed 100*μ*L of water to 100*μ*L of *Ct* culture (in MM 2×, OD = 0.2)) in addition to *Ct* in NC medium (in 4 wells we mixed 100*μ*L of water to 100*μ*L of *Ct* culture (in NC 2×, OD = 0.2)). For *hypoxanthine*, we proceeded slightly differently as its solubility is a lot lower than for the two other metabolites. We prepared a hypoxanthine stock at 5mM and directly added 200*μ*L to wells B-E(1). We then followed the same logic as for oxoglutarate and proline. We could test concentrations from 2.5mM to 0.002mM. The growth was assessed by measuring the OD_600_ every 10 minutes for 72 hours using a microplate reader (Biotek synergy H1, 28°C, continuous double-orbital shaking). The effect of each metabolites on the lag phase of *Ct* and on its final yield compared to both parameters in MM (positive controls) was assessed statistically, see tables S1, S2, S3, S4, S5, S6 (R version 4.1.2).

### Testing oxoglutarate, proline and hypoxanthine as carbon sources (fig.3D)

To test if the metabolites alone could support the growth of *Ct*, we grew *Ct* in NC medium containing intermediate concentrations of the metabolites (oxoglutarate and proline: 1.56mM; hypoxanthine: 1.25mM). We prepared *Ct* cultures (following the method described in “Cell culture preparations”) with an OD of 0.2 in water and in a 96-well plate mixed 100*μ*L of culture to 100*μ*L NC + metabolite 2×. The growth was assessed by measuring the OD_600_ every 10 minutes for 72 hours using a microplate reader (Biotek synergy H1, 28°C, continuous double-orbital shaking).

### Testing oxoglutarate, proline and hypoxanthine with increasing concentration of NC compounds on Ct

To test the effect of the metabolites on *Ct* when increasing concentrations of NC compounds are added, we chose an intermediate concentration of each metabolites (oxoglutarate and proline: 1.56mM; hypoxanthine: 1.25mM). We tested the growth of *Ct* in MM + metabolite when adding NC medium 2×, 1.5× or 1×. We thus prepared *Ct* cultures in MM + metabolites 2X with an OD of 0.2 and one culture of *Ct* in MM 2X only (as control). In a 96 well-plate, we added 100*μ*L of *Ct* culture to 100*μ*L of either NC 2×, 1.5×, 1× or water (as control). This way the MM is 1× concentrated and *Ct* is at an OD of 0.1 (as usually tested). As a negative control we also grew *Ct* in NC medium (OD = 0.1). The growth was assessed by measuring the OD_600_ every 10 minutes for 72 hours using a microplate reader (Biotek synergy H1, 28°C, continuous double-orbital shaking).

### Mathematical model

The mathematical model is described in supplementary note S3.

## ACKNOWLEDGEMENTS

We thank Christoph Keel, Rizlan Bernier-Latmani, Björn Vessman, Shota Shibasaki, Aurore Picot and Afra Salazar for very useful and constructive feedback on the manuscript. Untargeted metabolomic profiling was performed at the Metabolomics Platform, Faculty of Biology and Medicine, University of Lausanne. We acknowledge the entire team for their work from sample preparation and data acquisition to data processing and metabolite identification. We particularly thank Julijana Ivanisevic and Hector Gallart-Ayala for discussing the analyses with us. We also thank Alice Wallef, Nastassia Quévit for additional experiments, and Gwenaël Labouebe for access to the osmometer. We thank Björn Vessman for developing the first version of the mathematical model code https://gitlab.com/eccemic/facilitation2019. A.R.D.S. is funded by Swiss National Science Foundation grant PCEGP3_181272, R.D.M. by European Research Council grant 715097, S. T. by the NCCR Microbiomes, and S.M. by all three grants.

## Notes

### Competing Interest Statement

The authors have declared no competing interest.

