## Supplementary material for "Classifying interactions in a synthetic bacterial community is hindered by inhibitory growth medium"

### Supplementary Note S1: Supplementary figures and tables

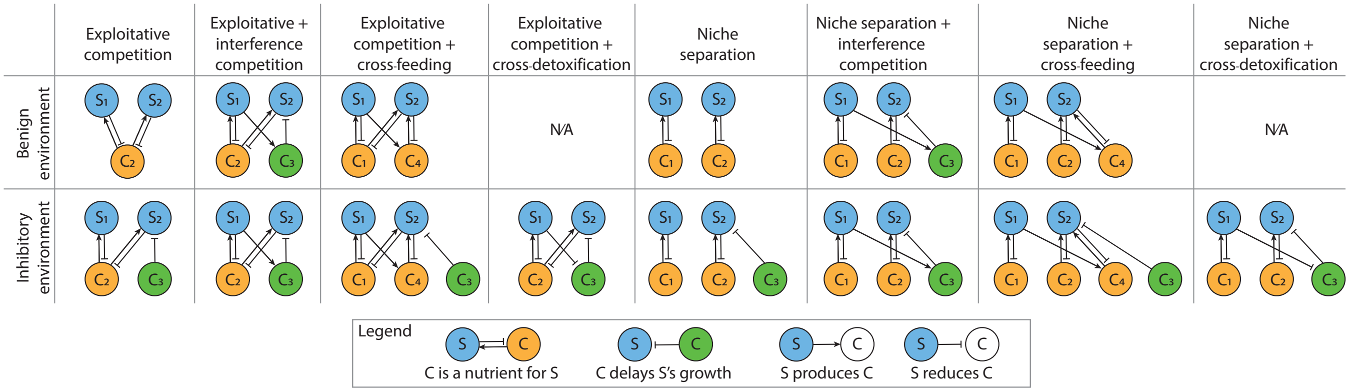


**Fig. S1.** Interactions used to build the model, which is described in Supplementary Note **??**. Note that interference competition looks the same in both environments, as our model assumes that it is mediated by the same compound $C_3$ that inhibits $S_2$'s growth in the inhibitory environment (in the inhibitory environment its concentration will be higher). In reality, of course, it is likely that a different inhibitory compound will be produced by $S_1$.

Model predictions under a benign environment are shown in Fig. S2, while those under the inhibitory environment, where $C_3$ increases $S_2$'s lag phase are shown in Fig. S6. Note that the only way to distinguish between cross-feeding (NS+CF) and cross-detoxification (NS+CD) is in the abundance of $S_2$ at stationary phase in condition IV relative to II.

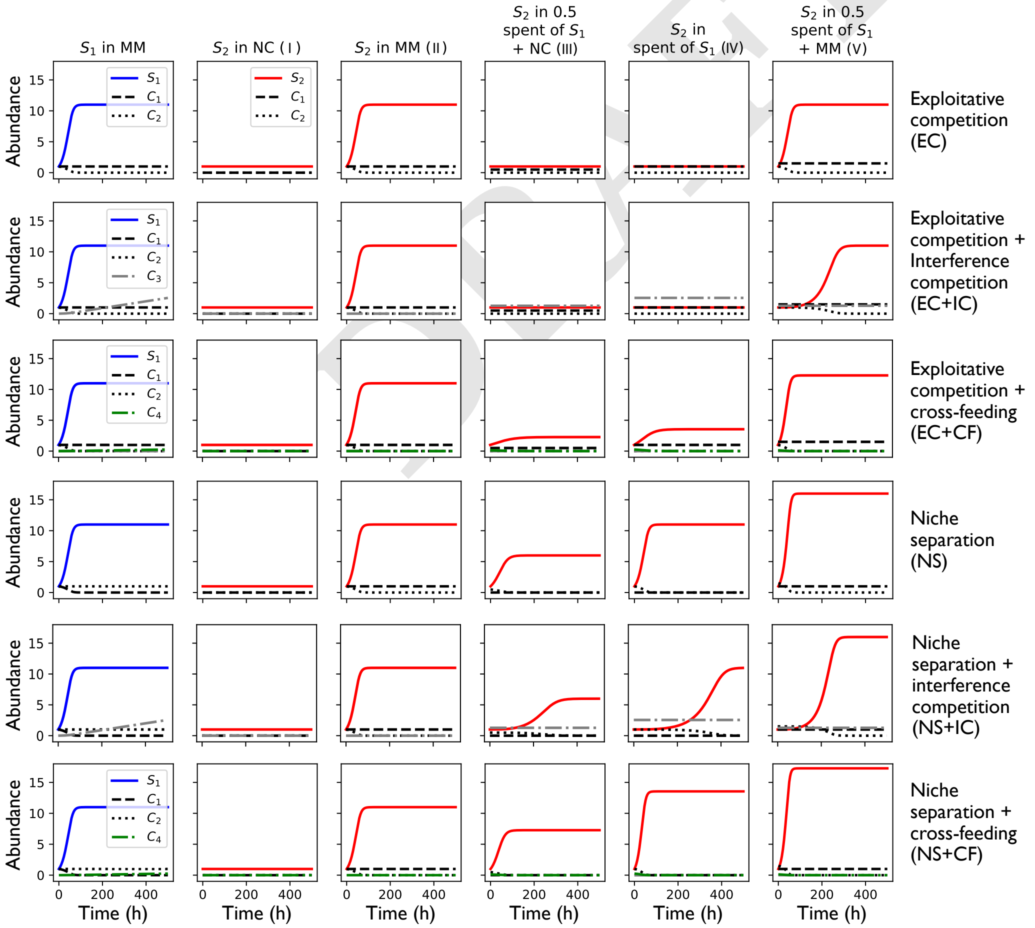


**Fig. S2.** Growth curves predicted by the model under a benign environment. Cross-detoxification does not apply here.

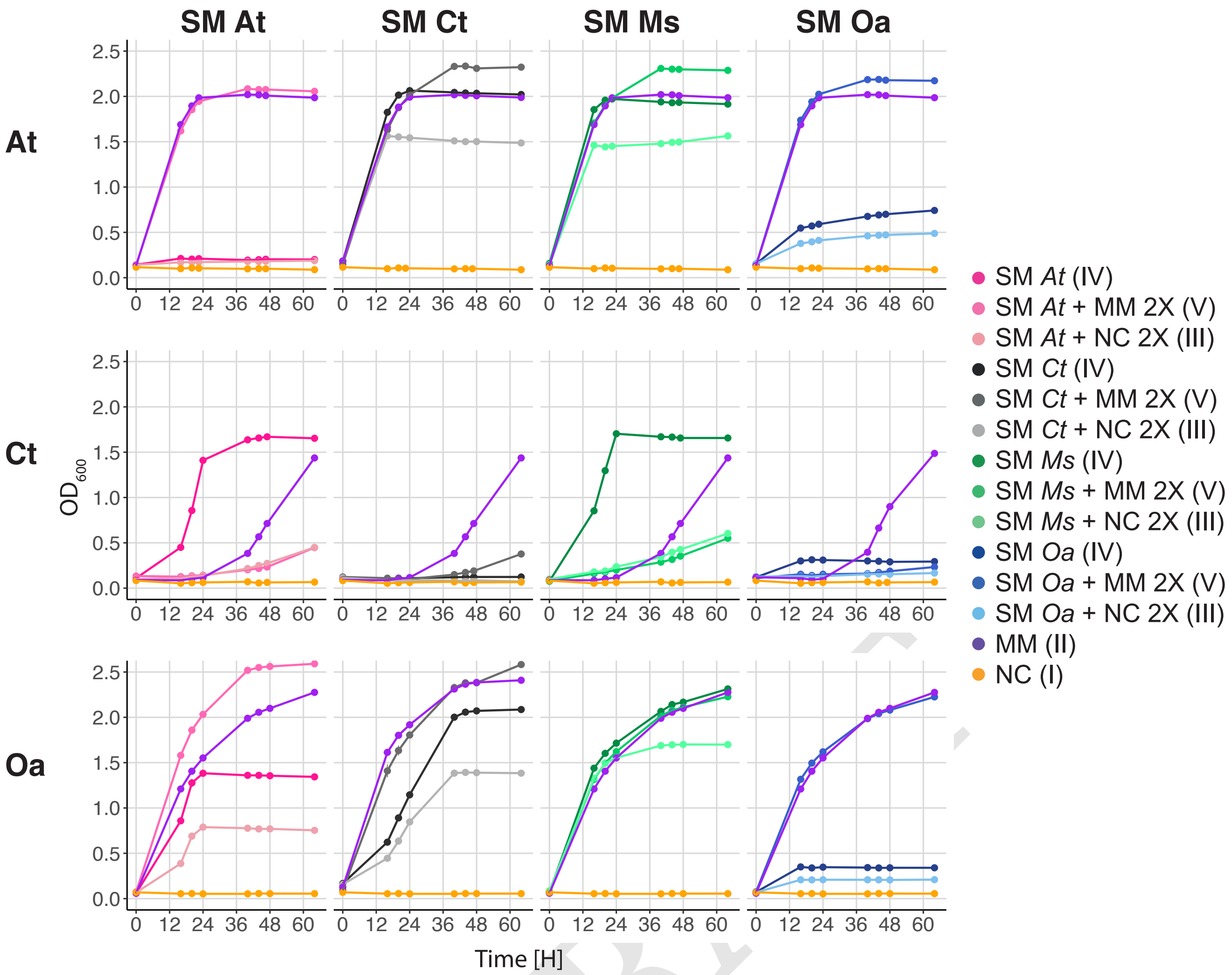


**Fig. S3.** $OD_{600}$ growth curves (raw data) from which AUCs were calculated in Fig. S4 (in parentheses the condition number is indicated similarly for each SM). Columns indicate the species that produced the spent medium (SM) and rows indicate which species is growing.

| Oxoglutarate | | | |
|---|---|---|---|
| **Concentration [mM]** | **p-values** | **Corrected p-values** | **Significance** |
| 12.5 | 1.52E-07 | 1.36E-06 | *** |
| 6.25 | 2.21E-07 | 1.99E-06 | *** |
| 3.12 | 1.57E-06 | 1.41E-05 | *** |
| 1.56 | 3.81E-07 | 3.42E-06 | *** |
| 0.78 | 8.32E-05 | 7.49E-04 | *** |
| 0.39 | 0.00100433 | 0.00903901 | ** |
| 0.19 | 0.06549582 | 0.589462338 | NS |
| 0.09 | 0.52441907 | 1 | NS |
| 0.04 | 0.87065699 | 1 | NS |

**Table S1.** Comparisons of the length of *Ct*'s lag phase in the different oxoglutarate concentrations compared to the length of *Ct*'s lag phase in MM (positive control) (see Fig. 3B). We estimated that *Ct*'s lag phase ended when the $OD_{600}$ exceeded 0.15 (visual estimation for this experiment). Here, oxoglutarate has an inhibitory effect on *Ct* at 50 and 25mM, and thus the $OD_{600}$ never exceeds 0.15 in those conditions. Multiple t-tests were performed (Welch two sample t-tests with Bonferroni correction) to evaluate if the length of the lag phase was smaller in presence of oxoglutarate.(* : p-value $<$ 0.05; ** : p-value $<$ 0.01; *** : p-value $<$ 0.001)

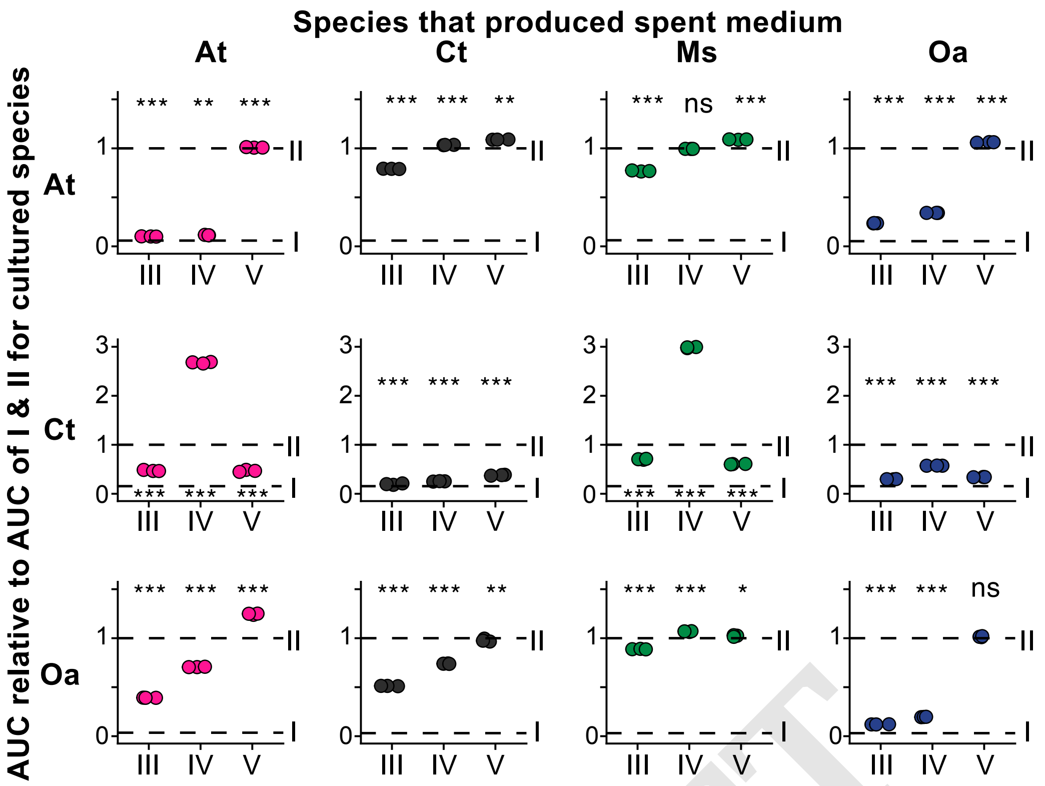


**Fig. S4.** SM assays on *At*, *Ct* and *Ms*. We cultured each species in five conditions: I. non-carbonic medium containing only salts and trace metals, but no carbon sources, II. medium I but with 15 mM glucose and 10 mM citric acid (CS: carbon sources), III. a mix of the spent medium of a given partner species (50%) and medium I (50% of a 2× concentrated solution), IV. the spent medium of a partner species, V. a mix of the spent medium of a partner species (50%) and medium II (50% of a 2× concentrated solution). The $OD_{600}$ was measured over time (see Fig. S3) and the area under the $OD_{600}$ growth curves was calculated. The AUC in each condition relatively to the AUC in II (dashed line II = 1) was plotted. The AUC in conditions III, IV, and V was compared to the AUC in II using multiple t-tests (two-sample equal variance t-tests) and Bonferroni corrections (* : p-value $<$ 0.05; ** : p-value $<$ 0.01; *** : p-value $<$ 0.001).

DRAFT

| **Proline** | | | |
|---|---|---|---|
| **Concentration [mM]** | **p-values** | **Corrected p-values** | **Significance** |
| 50 | 5.76E-04 | 0.006332828 | ** |
| 25 | 6.33E-04 | 0.006959028 | ** |
| 12.5 | 7.13E-04 | 0.007843465 | ** |
| 6.25 | 7.79E-04 | 0.008568448 | ** |
| 3.12 | 8.72E-04 | 0.00958818 | ** |
| 1.56 | 9.16E-04 | 0.010080192 | * |
| 0.78 | 0.00161288 | 0.017741693 | * |
| 0.39 | 0.00434241 | 0.047766496 | * |
| 0.19 | 0.00890951 | 0.098004556 | NS |
| 0.09 | 0.00718318 | 0.079015023 | NS |
| 0.04 | 0.03001905 | 0.330209514 | NS |

**Table S2.** Comparisons of the length of *Ct*'s lag phase in the different proline concentrations compared to the length of *Ct*'s lag phase in MM (positive control) (see Fig. 3B). We estimated that *Ct*'s lag phase ended when the $OD_{600}$ exceeded 0.13 (visual estimation for this experiment). Multiple t-tests were performed (Welch two sample t-tests with Bonferroni correction) to evaluate if the length of the lag phase was smaller in presence of proline.(* : p-value $<$ 0.05; ** : p-value $<$ 0.01; *** : p-value $<$ 0.001)

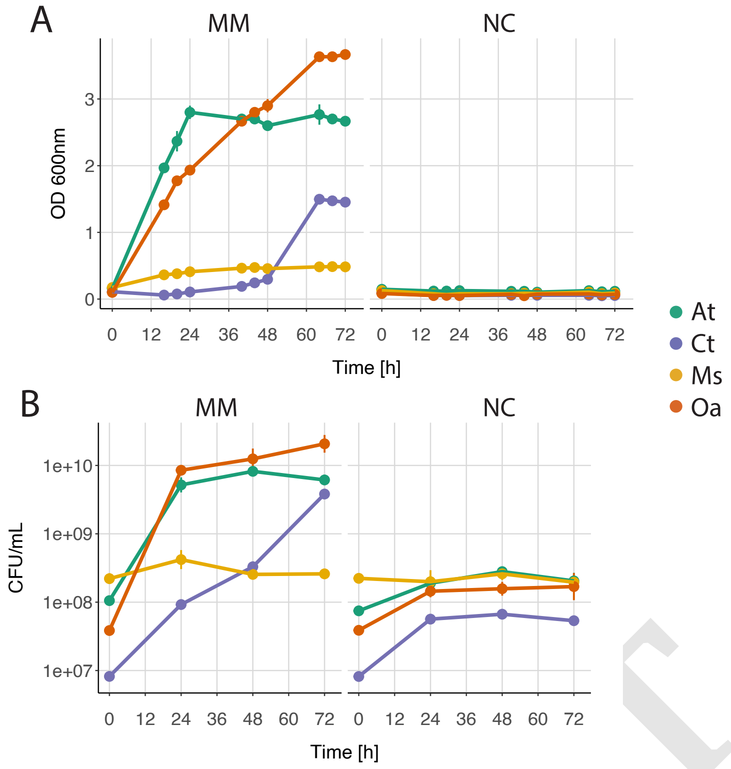


**Fig. S5.** Growth of *At*,*Ct*, *Ms* and *Oa* in monoculture in minimal medium (MM) and in no-carbon (NC) medium over 72 hours. (A): $OD_{600}$ ; (B): CFUs ($\pm$ sd), n= 3. Note that the $OD_{600}$ of *Ms* is increasing compared to the NC control, but not its CFU/mL. This led us to omit this species from our analysis.

DRAFT

| **Hypoxanthine** | | | |
|---|---|---|---|
| **Concentration [mM]** | **p-values** | **Corrected p-values** | **Significance** |
| 2.5 | 6.48E-04 | 0.007125092 | ** |
| 1.25 | 4.80E-04 | 0.005277212 | ** |
| 0.62 | 0.001836386 | 0.020200247 | * |
| 0.31 | 0.004750006 | 0.05225007 | NS |
| 0.15 | 0.019130465 | 0.210435118 | NS |
| 0.078 | 0.108759136 | 1 | NS |
| 0.039 | 0.117378975 | 1 | NS |
| 0.019 | 0.193505308 | 1 | NS |
| 0.0097 | 0.287559421 | 1 | NS |
| 0.0048 | 0.091108658 | 1 | NS |
| 0.0024 | 0.183978005 | 1 | NS |

**Table S3.** Comparisons of the length of *Ct*'s lag phase in the different hypoxanthine concentrations compared to the length of *Ct*'s lag phase in MM (positive control) (see Fig. 3B). We estimated that *Ct*'s lag phase ended when the $OD_{600}$ exceeded 0.13 (visual estimation for this experiment). Multiple t-tests were performed (Welch two sample t-tests with Bonferroni correction) to evaluate if the length of the lag phase was smaller in presence of hypoxanthine.(* : p-value $<$ 0.05; ** : p-value $<$ 0.01; *** : p-value $<$ 0.001)

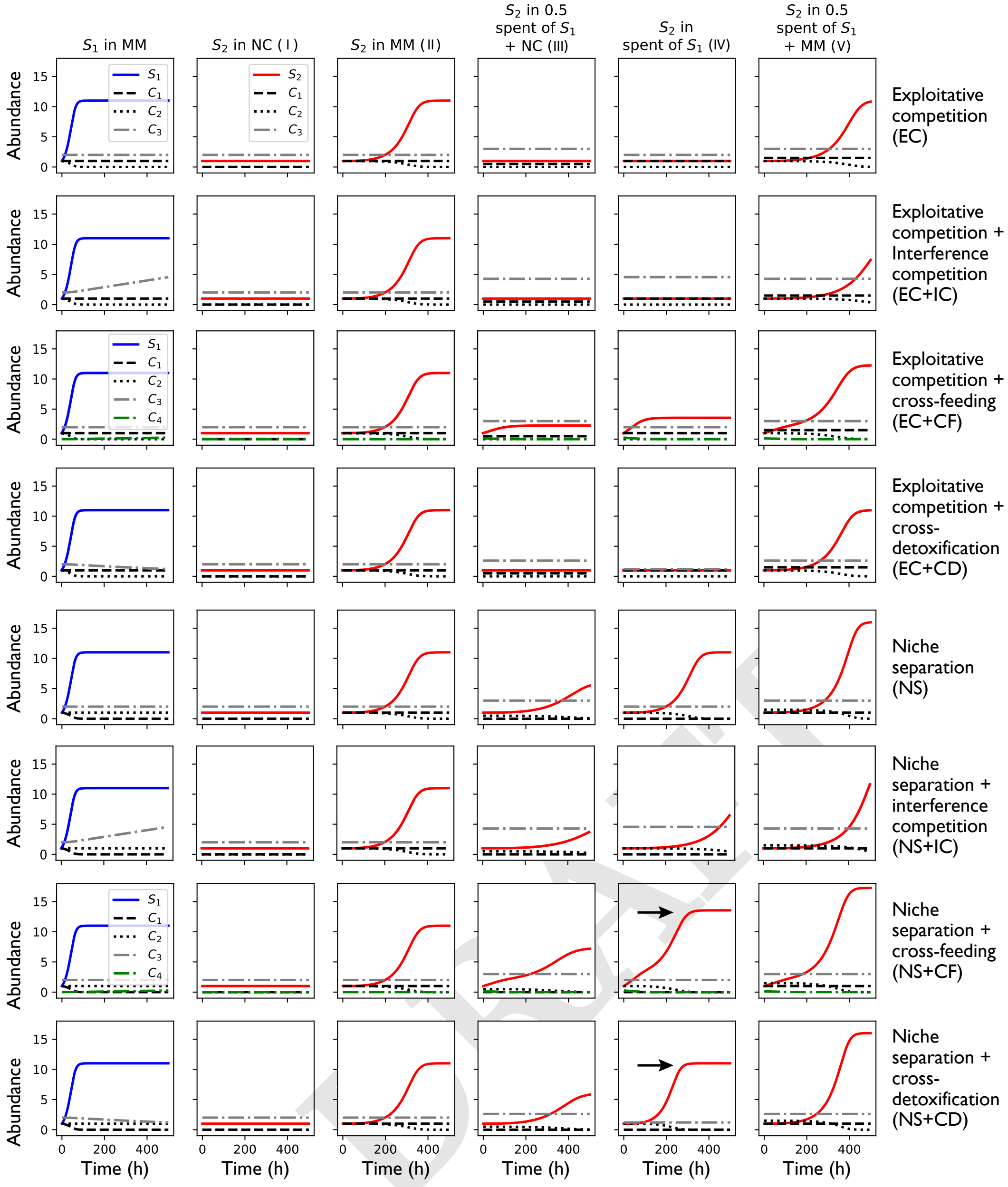


**Fig. S6.** Growth curves predicted by the model under an inhibitory environment.

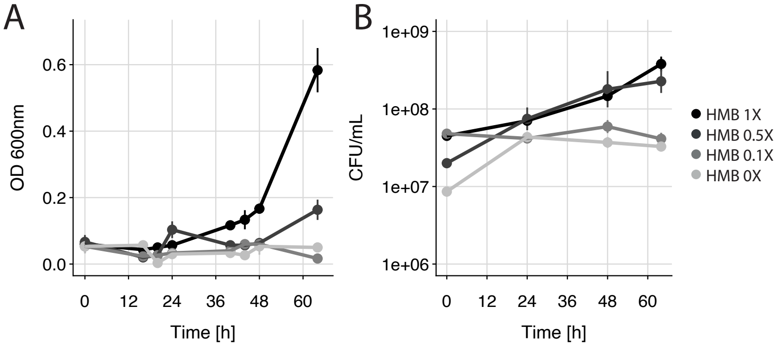


**Fig. S7.** Growth of *Ct* in MM with variations of HMB concentration. (A): Growth as $OD_{600}$ over time. (B): Growth as CFUs over time. Decresing the concentration of HMB is detrimental for *Ct*'s growth. (mean is plotted ±sd, n = 3)

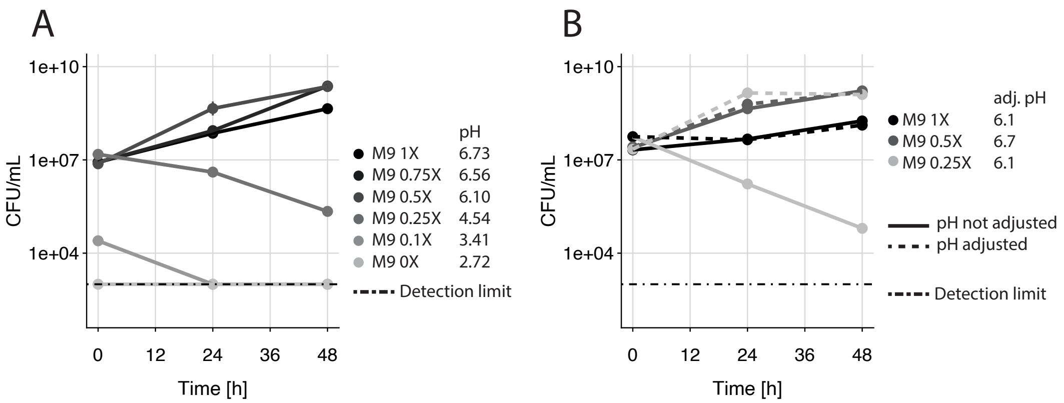


**Fig. S8.** (A) Growth (CFUs over time, see Fig. 2A for $OD_{600}$) in MM with decreasing concentrations of M9. The resulting pH is indicated for each fresh MM, showing that decreasing the concentration of M9 also decreases the pH, which impairs *Ct*'s growth. (B) Growth (CFUs over time, see Fig. 2B for $OD_{600}$) in MM with decreasing concentrations of M9 with and without manual pH adjustment to either 6.1 or 6.7 using NaOH. Decreasing the concentration of M9 while keeping a pH close to neutrality shortens the lag phase of *Ct*.(Plotted: mean $\pm$sd, n = 3)

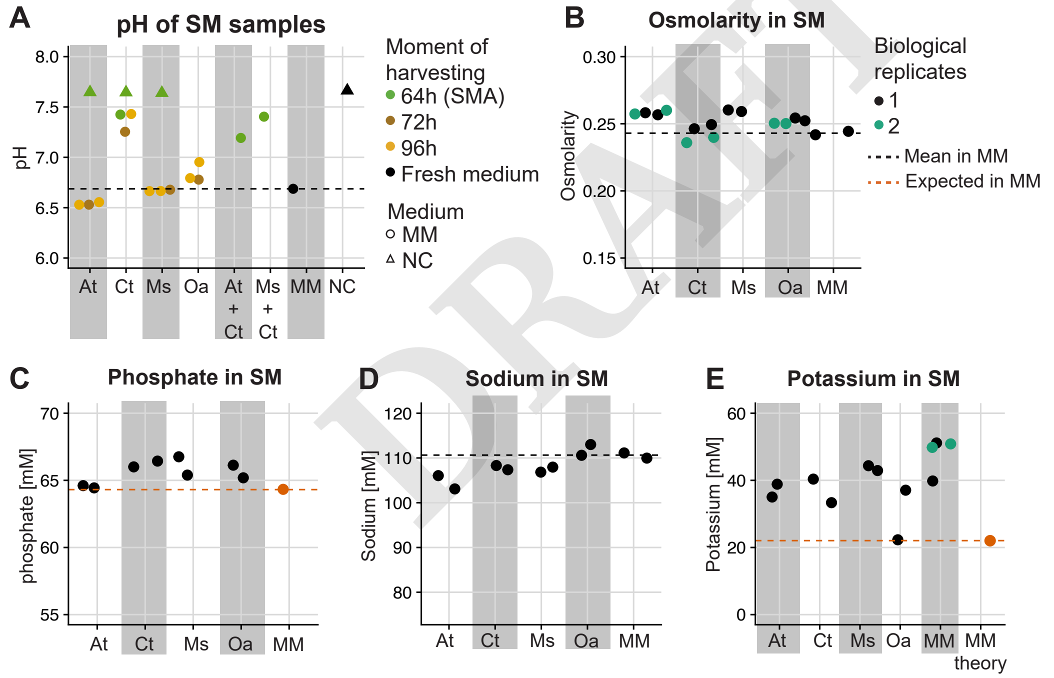


**Fig. S9.** Chemical analysis of the four species' spent media (SM). (A) pH of SM of *At*, *Ct*, *Ms* and *Oa* in stationary phase (72 or 96 hours of growth), and SM of *Ct* grown in SM of *At* and *Ms* after 64 hours (length of initial SM assay, abbreviated SMA). As controls we measured the pH of the fresh minimal medium (MM=NC+CS, black dashed line) and the no-carbon salts and trace metals (NC). Panels (B)-(E) show (B) osmolarity, (C) Phosphate quantification, (D) sodium quantification and (E) potassium quantification. Data in panels (C)-(E) come from commercial chemical kits. Black dashed lines indicate read-outs from MM (A, B and D)). The orange dashed line in (C) and (E) represents the theoretical potassium and phosphate,respectively, concentration in MM. NB: no statistics were performed as all measures were done on technical duplicates (except for panel B were we have biological duplicates in addition to technical duplicates). We conclude from these analyses that *At* and Ms do not modify the chemical environment in ways that should shorten the lag phase of *Ct*.

DRAFT

A

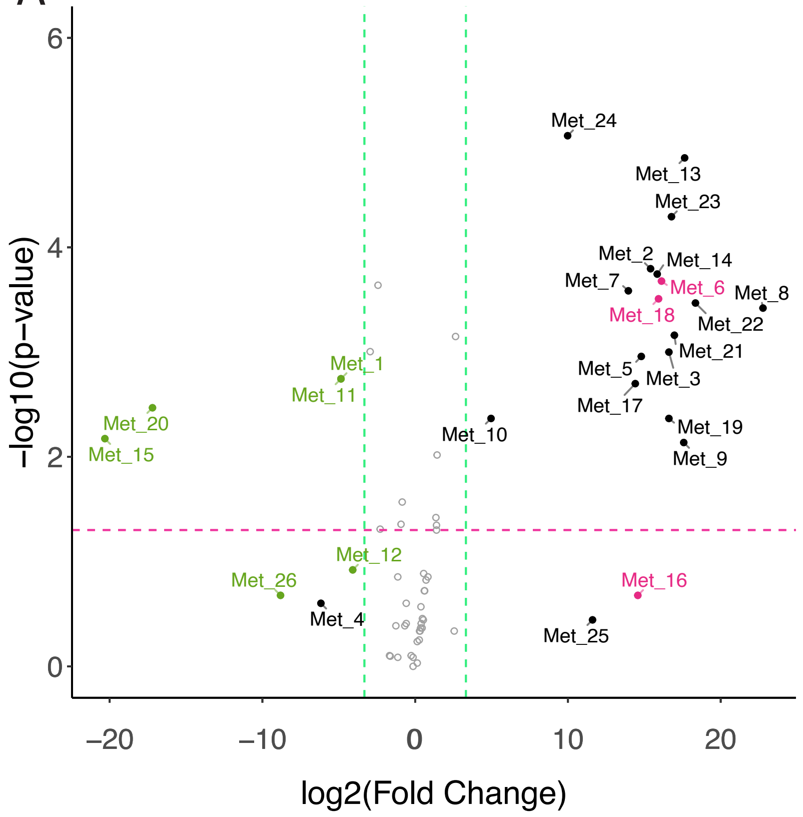


| Metabolite | Number |
|---|---|
| 2_5-DIHYDROXYBENZOATE | Met_1 |
| 4-GUANIDINOBUTANOATE | Met_2 |
| 6-HYDROXYNICOTINATE | Met_3 |
| ALPHA-D-GLUCOSE 2 | Met_4 |
| BETA-NICOTINAMIDE_ADENINE_DINUCLEOTIDE | Met_5 |
| CITRULLINE | Met_6 |
| CYTIDINE | Met_7 |
| DEOXYADENOSINE | Met_8 |
| DEOXYCYTIDINE | Met_9 |
| DEOXYGUANOSINE | Met_10 |
| DIHYDROXYBENZOATE_1 | Met_11 |
| GALACTOSAMINE | Met_12 |
| GUANOSINE | Met_13 |
| GUANOSINE_MONOPHOSPHATE | Met_14 |
| HYPOXANTHINE | Met_15 |
| MELIBIOSE | Met_16 |
| N-ACETYLGALACTOSAMINE | Met_17 |
| N_N-DIMETHYLARGININE | Met_18 |
| PIPECOLATE | Met_19 |
| PTERIN | Met_20 |
| THYMIDINE-MONOPHOSPHATE | Met_21 |
| URIDINE | Met_22 |
| URIDINE_5-DIPHOSPHATE | Met_23 |
| URIDINE_DIPHOSPHATE-N-ACETYLGLUCOSAMINE | Met_24 |
| URIDINE_DIPHOSPHATE_GLUCOSE | Met_25 |
| XANTHINE | Met_26 |

B

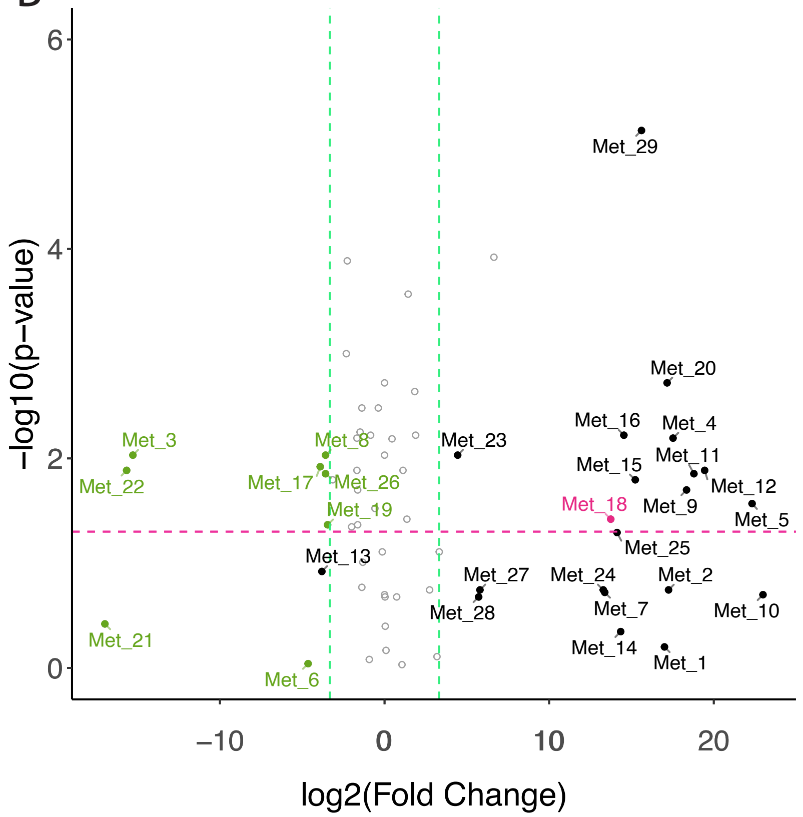


| Metabolite | Number |
|---|---|
| 4-ACETAMIDOBUTANOATE | Met_1 |
| 4-GUANIDINOBUTANOATE | Met_2 |
| 4-HYDROXY-L-PHENYLGLYCINE | Met_3 |
| 6-HYDROXYNICOTINATE | Met_4 |
| ADENINE | Met_5 |
| ARGININE | Met_6 |
| BETA-NICOTINAMIDE_ADENINE_DINUCLEOTIDE | Met_7 |
| CIS-4-HYDROXY-D-PROLINE | Met_8 |
| CYTIDINE | Met_9 |
| DEOXYADENOSINE | Met_10 |
| DEOXYCYTIDINE | Met_11 |
| DEOXYGUANOSINE | Met_12 |
| GALACTOSAMINE | Met_13 |
| GUANOSINE_MONOPHOSPHATE | Met_14 |
| N-ACETYLGALACTOSAMINE | Met_15 |
| N-ACETYLPUTRESCINE | Met_16 |
| N-ALPHA-ACETYLLYSINE | Met_17 |
| N_N-DIMETHYLARGININE | Met_18 |
| OXOGLUTARATE | Met_19 |
| PIPECOLATE | Met_20 |
| PROLINE | Met_21 |
| PTERIN | Met_22 |
| PYRIDOXAMINE | Met_23 |
| S-ADENOSYLHOMOCYSTEINE | Met_24 |
| S-ADENOSYLMETHIONINE | Met_25 |
| TRANS-4-HYDROXY-L-PROLINE | Met_26 |
| URACIL | Met_27 |
| URIDINE | Met_28 |
| URIDINE_5-DIPHOSPHATE | Met_29 |

**Fig. S10.** Identification of metabolites consumed and produced by *Ct* in the spent media of *At* (A) and *Ms* (B) (same data as seen on Fig. 3A). Tables indicate the full names of the metabolites that had a fold-change of at least 10.

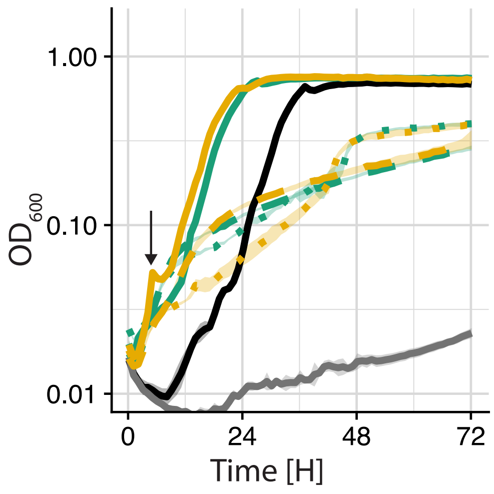


**Fig. S11.** Growth curves of Fig. 4D with the Y axis in $\log 10$.

DRAFT

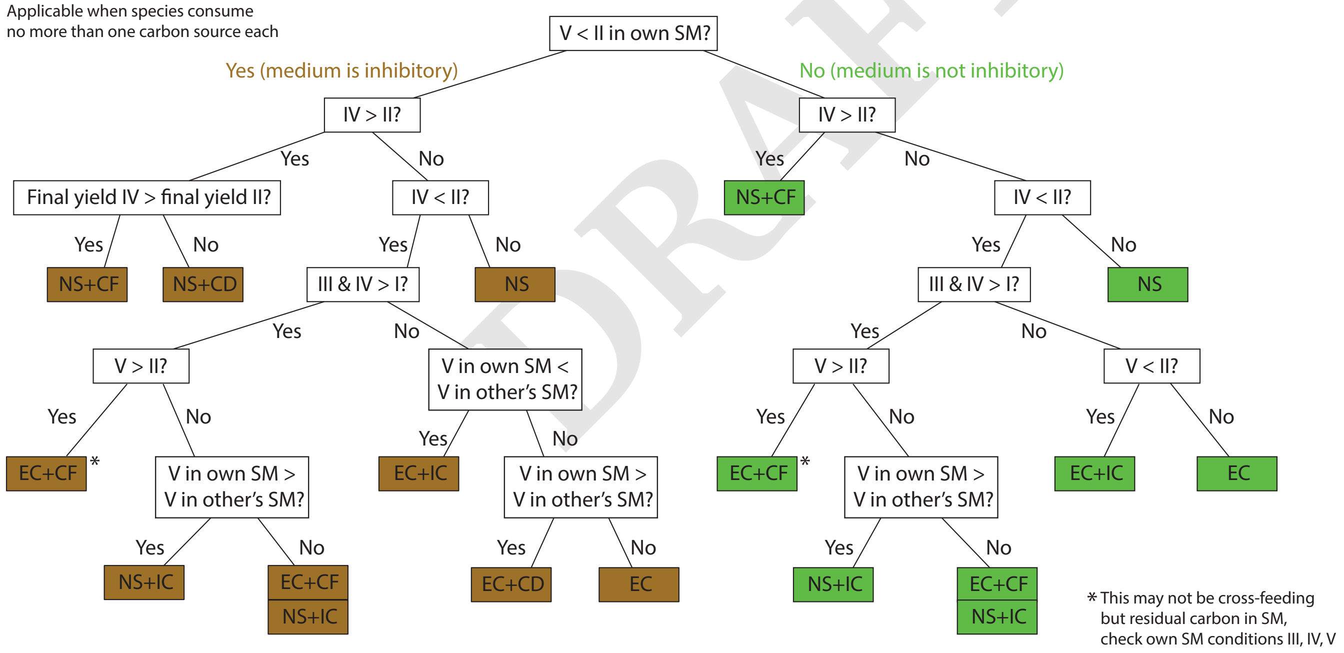


**Fig. S12.** Decision tree to help classify interactions. Colors indicate leaf nodes (interaction types) either under an inhibitory environment (brown) or a benign environment (green). Abbreviations are as in Fig. 4 and S6. The tree applies only when the species that produced the SM and the one whose growth is being analysed each consume a single carbon source. Two conditions are difficult to distinguish: EC+CF and NS+IC. The tree includes some hints of how they may be distinguished, but it is still possible that the results remain ambiguous. It is also worth noting that a result that looks like EC+CF may be due to residual carbon in the SM medium (that was not consumed entirely by the SM producer). This can be identified by analysing the SM producer growing in its own SM: did it also grow more in condition V than in condition II? If this is the case, it suggests residual carbon rather than cross-feeding.

| Oxoglutarate | | | |
|---|---|---|---|
| Concentration [mM] | p-values | Corrected p-values | Significance |
| 50 | 0.99999944 | 1 | NS |
| 25 | 1 | 1 | NS |
| 12.5 | 2.16E-08 | 2.38E-07 | *** |
| 6.25 | 1.18E-08 | 1.30E-07 | *** |
| 3.12 | 3.32E-06 | 3.65E-05 | *** |
| 1.56 | 0.00482831 | 0.053111395 | NS |
| 0.78 | 0.00357138 | 0.039285195 | * |
| 0.39 | 0.00376815 | 0.041449632 | * |
| 0.19 | 0.00141323 | 0.015545504 | * |
| 0.09 | 0.00403497 | 0.044384673 | * |
| 0.04 | 0.01703349 | 0.18736836 | NS |

**Table S4.** Comparisons of the final yield of *Ct* in the different oxoglutarate concentrations tested compared to its final yield in MM (positive control) (see Fig. 3B). Multiple t-tests were performed (Welch two sample t-tests with Bonferroni correction) to evaluate if the final yield was higher in presence of oxoglutarate.(* : p-value $<$ 0.05; ** : p-value $<$ 0.01; *** : p-value $<$ 0.001)

DRAFT

| Proline | | | |
|---|---|---|---|
| Concentration [mM] | p-values | Corrected p-values | Significance |
| 50 | 9.94E-07 | 1.09E-05 | *** |
| 25 | 3.32E-11 | 3.65E-10 | *** |
| 12.5 | 1.00E-06 | 1.10E-05 | *** |
| 6.25 | 1.70E-07 | 1.87E-06 | *** |
| 3.12 | 6.74E-06 | 7.41E-05 | *** |
| 1.56 | 0.001611674 | 0.017728416 | * |
| 0.78 | 0.002418454 | 0.026602993 | * |
| 0.39 | 0.023539314 | 0.258932458 | NS |
| 0.19 | 0.032234754 | 0.354582295 | NS |
| 0.09 | 0.165837609 | 1 | NS |
| 0.04 | 0.842998364 | 1 | NS |

**Table S5.** Comparisons of the final yield of *Ct* in the different proline concentrations tested compared to its final yield in MM (positive control) (see Fig. 3B). Multiple t-tests were performed (Welch two sample t-tests with Bonferroni correction) to evaluate if the final yield was higher in presence of proline.(* : p-value $<$ 0.05; ** : p-value $<$ 0.01; *** : p-value $<$ 0.001)

| Hypoxanthine | | | |
|---|---|---|---|
| Concentration [mM] | p-values | Corrected p-values | Significance |
| 2.5 | 2.11E-04 | 0.002322673 | ** |
| 1.25 | 3.73E-04 | 0.004106511 | ** |
| 0.62 | 0.703957543 | 1 | NS |
| 0.31 | 0.916597085 | 1 | NS |
| 0.15 | 0.303073963 | 1 | NS |
| 0.078 | 0.958711043 | 1 | NS |
| 0.039 | 0.996391847 | 1 | NS |
| 0.019 | 0.730097349 | 1 | NS |
| 0.0097 | 0.906572689 | 1 | NS |
| 0.0048 | 0.234967922 | 1 | NS |
| 0.0024 | 0.885431801 | 1 | NS |

**Table S6.** Comparisons of the final yield of *Ct* in the different hypoxanthine concentrations tested compared to its final yield in MM (positive control) (see Fig. 3B). Multiple t-tests were performed (Welch two sample t-tests with Bonferroni correction) to evaluate if the final yield was higher in presence of hypoxanthine.(* : p-value $<$ 0.05; ** : p-value $<$ 0.01; *** : p-value $<$ 0.001)

| Ct in the SM of At and Ms - Comparison of III-V to II | | | |
|---|---|---|---|
| Condition | p-values | corrected p-values | Significance |
| SM At - III | 0.99999999 | 1 | NS |
| SM Ms - III | 0.999999695 | 1 | NS |
| SM At - IV | 2.14E-04 | 0.001286046 | ** |
| SM Ms - IV | 3.19E-07 | 1.92E-06 | *** |
| SM At - V | 0.999999999 | 1 | NS |
| SM Ms - V | 0.999977537 | 1 | NS |

**Table S7.** Comparison of the final yield of *Ct* in conditions III, IV and V compared to condition II (MM, positive control) (see Fig. 1A for detailed description of these conditions and Fig. 4D for the growth curves). Multiple t-tests were performed (Welch two sample t-tests with Bonferroni correction) to evaluate if the final yield was higher in the tested conditions than in condition II.(* : p-value $<$ 0.05; ** : p-value $<$ 0.01; *** : p-value $<$ 0.001)

DRAFT

| Ct in the SM of At and Ms - Comparison of III-V to II | | | |
|---|---|---|---|
| Condition | p-values | corrected p-values | Significance |
| SM At - III | 8.86E-08 | 5.31E-07 | *** |
| SM Ms - III | 3.64E-07 | 2.19E-06 | *** |
| SM At - IV | 6.38E-07 | 3.83E-06 | *** |
| SM Ms - IV | 1.82E-07 | 1.09E-06 | *** |
| SM At - V | 1.90E-08 | 1.14E-07 | *** |
| SM Ms - V | 2.63E-09 | 1.58E-08 | *** |

**Table S8.** Comparison of the length of *Ct*'s lag phase in conditions III, IV and V compared to condition II (MM, positive control) (see Fig. 1A for detailed description of these conditions and Fig. 4D for the growth curves). We estimated that *Ct*'s lag phase ended when the $OD_{600}$ exceeded 0.04 (visual estimation for this experiment). Multiple t-tests were performed (Welch two sample t-tests with Bonferroni correction) to evaluate if the length of the lag phase was smaller in the tested conditions than in condition II. It should be pointed out that in conditions III and V (Fig. 4D) *Ct* grows slowly but gradually and exceeds an $OD_{600}$ of 0.04 early, but without entering an exponential phase. Therefore, focusing on the shapes of the growth curves, we considered that only in condition IV the lag phase of *Ct* is actually shorter that in condition II. (* : p-value $<$ 0.05; ** : p-value $<$ 0.01; *** : p-value $<$ 0.001)

| Solution | Volume [mL] |
|---|---|
| M9 10× | 100 |
| HMB 50× | 20 |
| Glucose 0.75M | 20 |
| Citric acid 0.5M | 20 |
| Water | 840 |

**Table S9.** Minimal medium (MM) with glucose and citric acid (MM) - 1L

| Solution | Volume [mL] |
|---|---|
| M9 10× | 100 |
| HMB 50X× | 20 |
| Water | 880 |

**Table S10.** No-Carbon (NC) medium - 1L

| Compound | Quantity |
|---|---|
| NTA | 10 g |
| $MgSO_4 \times 7H_2O$ | 14.45 g |
| $CaCl_2 \times 2H_2O$ | 3.33 g |
| $(NH_4)_6Mo_7O_{24} \times 4H_2O$ | 0.00974 g |
| $FeSO_4 \times 7H_2O$ | 0.099 g |
| Metals 44 | 50 mL |
| H2O | up to 1 L |

**Table S11.** Composition of Hutner's vitamin-free Mineral Base (HMB) 50× [1L].

| Compound | Mass [g] |
|---|---|
| $Na_2EDTA$ x $2H_2O$ | 0.387 |
| $ZnSO_4$ x $7H_2O$ | 1.095 |
| $FeSO_4$ x $7H_2O$ | 0.914 |
| $MnSO_4$ x $H_2O$ | 0.154 |
| $CuSO_4$ x $5H_2O$ | 0.0392 |
| $Co(NO3)_2$ x $6H_2O$ | 0.0248 |
| $Na_2B_4O_7$ x $10H_2O$ | 0.0177 |
| H2O | up to 100 mL |

**Table S12.** Composition of the solution Metal 44 used in the HMB

| Powder | Mass [g] |
|---|---|
| $Na_2HPO_4$ | 60 |
| $KH_2PO_4$ | 30 |
| NaCl | 5 |
| $NH_4Cl$ | 10 |

**Table S13.** Composition of M9 10× [1L]. Mix all ingredients in 800 mL of distilled water, adjust the pH to 7.4 with NaOH, add water up to 1L. Autoclave.

DRAFT

| Powder | Mass [g] |
|---|---|
| $Na_2HPO_4$ | 72.40 |
| $KH_2PO_4$ | 17.70 |
| NaCl | 5 |
| $NH_4Cl$ | 10 |

**Table S14.** Composition of M9 10× [1L] without pH adjustment needed. Mix all ingredients in 1L of distilled water. Autoclave. Final pH ≅ 7.4

| Powder | Mass [g] |
|---|---|
| $Na_2HPO_4$ | 72.40 |
| $NaH_2PO_4$ | 15.60 |
| NaCl | 5 |
| $NH_4Cl$ | 10 |

**Table S15.** Composition of M9 10× [1L] with only sodium ions in the phosphate buffer. Mix all ingredients in 1L of distilled water. Autoclave.Final pH ≅ 7.4

| Powder | Mass [g] |
|---|---|
| $K_2HPO_4$ | 88.80 |
| $KH_2PO_4$ | 17.70 |
| NaCl | 5 |
| $NH_4Cl$ | 10 |

**Table S16.** Composition of M9 10× [1L] with only potassium ions in the phosphate buffer. Mix all ingredients in 1L of distilled water. Autoclave. Final pH ≅ 7.4

DRAFT

### Supplementary Note S2: Metabolomics analyses

**Metabolite extraction:** To extract the metabolites, 400 $\mu$L of ice-cold MeOH was added to 100 $\mu$L of each medium. The samples were then vortexed for 30 seconds, followed by 15 min centrifugation at 13,000 rpm at 4° C. The resulting supernatant was collected and evaporated to dryness in a vacuum concentrator (LabConco, Missouri, US) and the dried extract was reconstituted in 100 $\mu$L of 80 % MeOH prior to LC-MS injection (61). **Data acquisition - LC-HRMS analyses:** Media extracts were analyzed by Hydrophilic Interaction Liquid Chromatography coupled to high resolution mass spectrometry (HILIC - HRMS) in both positive and negative ionization modes using a 6550 Quadrupole Time-of-Flight (Q-TOF) system interfaced with 1290 UHPLC system (Agilent Technologies) as previously described (62). In positive mode, the chromatographic separation was carried out in an Acquity BEH Amide, 1.7 $\mu$m, 100 mm × 2.1 mm I.D. column (Waters, Massachusetts, US). Mobile phase was composed of A = 20 mM ammonium formate and 0.1 % FA in water and B = 0.1 % formic acid in ACN. The linear gradient elution from 95% B (0-1.5 min) down to 45% B was applied (1.5 min -17 min) and this conditions were held for 2 min. Then initial chromatographic condition were maintained as a post-run during 5 min for column re-equilibration. The flow rate was 400 $\mu$L/min, column temperature 25 °C and sample injection volume 2$\mu$l. In negative mode, a SeQuant ZIC-pHILIC (100 mm, 2.1 mm I.D. and 5 $\mu$m particle size, Merck, Damstadt, Germany) column was used. The mobile phase was composed of A = 20 mM ammonium Acetate and 20 mM NH4OH in water at pH 9.7 and B = 100% ACN. The linear gradient elution from 90% (0-1.5 min) to 50% B (8-11 min) down to 45% B (12-15 min). Finally, the initial chromatographic conditions were established as a post-run during 9 min for column re-equilibration (63). The flow rate was 300 $\mu$L/min, column temperature 30 °C and sample injection volume 2$\mu$l. Mass spectrometry ESI source conditions were set as follows: dry gas temperature 290 °C and flow 14 L min-1, fragmentor voltage 380 V, sheath gas temperature 350 °C and flow 12 L min-1, nozzle voltage 0 V, and capillary voltage +2000 V in positive mode and -2000 V in negative ionization mode. The instrument was set to acquire over the full m/z range 50-1000 in both modes, with the MS acquisition rate of 2 spectra/s. In addition, AIF (all ion fragmentation) MS/MS analysis were performed on pooled QC samples at a collision energy (CE) of 0, 10 and 30 eV. **Quality control (QC):** Pooled QC samples (representative of the entire sample set) were analyzed periodically (every 6 samples) throughout the overall analytical run in order to assess the quality of the data, correct the signal intensity drift (attenuation in most cases, that is inherent to LC-MS technique and MS detector due to sample interaction with the instrument over time) and remove the peaks with poor reproducibility (CV $> 30\%$) (64). In addition, a series of diluted quality controls (dQC) were prepared by dilution with methanol: $100\%$ QC, $50\%$ QC, $25\%$ QC, $12.5\%$ QC and $6.25\%$ QC and analyzed at the beginning and at the end of the sample batch. This QC dilution series served as a linearity filter to remove the features which don't respond linearly or for which the correlation with dilution factor was $< 0.75$. **Data (pre) processing:** Raw LC/MS files were processed using Profinder B.08.00 software (Agilent Technologies) for metabolite identification using an in-house database containing around 600 metabolites. Metabolites were identified based on accurate mass and retention time matching against standards solutions characterized under the same LC-MS conditions and the parameters settings were as follows: Match tolerance masses 10 ppm, Retention time tolerance 0.2 min, height filter 1000 counts, peak spectrum obtained as an average of scants at $10\%$ of the peak. The relative quantification of metabolites was based on EIC (Extracted Ion Chromatogram) areas. The obtained tables (containing peak areas of detected metabolites across all samples) were exported to "R" software `http://cran.r-project.org/` and signal intensity drift correction was done within the LOWESS/Spline normalization program (63) followed by noise filtering (CV (QC features) $> 30\%$). **Metabolite identification:** Short listed ions of interest were matched against the in-house created Accurate Mass Retention Time (AMRT) database to confirm the metabolite identities. Putatively identified metabolite features were further subjected to fragmentation (MS/MS data) pattern matching. The metabolite identifications were validated by matching the deconvoluted MS/MS against METLIN standard metabolite database and in-house recorded spectral library acquired on standards (62). **Statistical analyses (univariate):** ANOVA one-factor (on $\log 10$ transformed data) was used to test the significance of metabolite changes in different conditions (i.e. different fresh and spent media) with an arbitrary level of significance, p-value $= 0.05$.

DRAFT

### Supplementary Note S3: Mathematical model

Consider two species that each grow alone in a well-mixed culture medium containing several chemical compounds. The concentration $C_j$ of each compound $j$ can affect the abundance $S_i$ of each species $i \in \{1,2\}$ via feeding or growth retardation. When compound $j$ is a nutrient, its effect on $S_i$ follows the growth function $\rho_i$ that saturates with increasing compound concentrations (Monod growth) and the concentration of compound $j$ decreases as a function of $S_i$ via the biomass yield $Y_{i,j}$ and growth function $\rho_i$. Our first version of the model considered only such positive effects of compounds on growth. In our second version of the model (see discussion section of the main text), we augmented our model such that a compound $k$ can possibly have a negative effect on $S_i$ by increasing its lag phase. Based on our experimental observations, the lag phase is increased only for growth on certain compounds $j$ when compound $k$ is present. This is captured by the second term in the growth function $\rho_i$ and modulated by lag factor $l_{i,j,k}$. Finally, compound $j$ can be produced by species $i$ at rate $p_{i,j}$, or passively taken up at rate $u_{i,j}$. This results in the following set of differential equations:

$$\frac{dS_i}{dt} = \sum_j \rho_i(C_j, C_k) S_i \quad \textbf{(1a)}$$

$$\rho_i(C_j, C_k) = r_{i,j} \frac{C_j}{C_j + K_{i,j}} \frac{t^n}{(\sum_{k \neq j} l_{i,j,k} C_k)^n + t^n} \quad \textbf{(1b)}$$

$$\frac{dC_j}{dt} = \begin{cases} -\frac{1}{Y_i}\rho_i(C_j, C_k) S_i & \text{if } C_j \text{ is a nutrient for species } i \\ p_{i,j} S_i & \text{if } C_j \text{ is produced by species } i \\ -u_{i,j} C_j S_i & \text{if } C_j \text{ is passively taken up by species } i \end{cases} \quad \textbf{(1c)}$$

$Y_{i,j}$ is always 0.1, $K_{i,j}$ is always 1.0. The exponent $n$ is 2 and $t$ refers to time. The different interaction types are illustrated in Fig. S1. Depending on the type of interaction modeled, parameters $r_{i,j}$, $p_{i,j}$ and $l_{i,j}$ can vary as indicated in Table S17 below. Values not listed there (e.g. $r_{2,1}$) are 0 throughout all our simulations. The model was implemented in Python 3.8.2 using the SciPy library v1.7.1. ODEs were solved using scipy.integrate.ode and integrator dopri5, which uses the Runge-Kutta method.

|  | EC | EC+IC | EC+CF | EC+CD | NS | NS+IC | NS+CF | NC+CD |
|---|---|---|---|---|---|---|---|---|
| $r_{1,1}$ | 0 | 0 | 0 | 0 | 0.1 | 0.1 | 0.1 | 0.1 |
| $r_{1,2}$ | 0.1 | 0.1 | 0.1 | 0.1 | 0 | 0 | 0 | 0 |
| $r_{2,2}$ | 0.1 | 0.1 | 0.1 | 0.1 | 0.1 | 0.1 | 0.1 | 0.1 |
| $r_{2,4}$ | 0 | 0 | 0.1 | 0 | 0 | 0 | 0.1 | 0 |
| $p_{1,3}$ | 0 | 0.1 | 0 | 0 | 0 | 0.1 | 0 | 0 |
| $p_{1,4}$ | 0 | 0 | $10^{-5}$ | 0 | 0 | 0 | $10^{-5}$ | 0 |
| $l_{2,2,3}$ | 300 | 300 | 300 | 300 | 300 | 300 | 300 | 300 |
| $u_{1,3}$ | 0 | 0 | 0 | 0.0001 | 0 | 0 | 0 | 0.0001 |
| $S_1(t=0)$ | 1.0 | 1.0 | 1.0 | 1.0 | 1.0 | 1.0 | 1.0 | 1.0 |
| $S_2(t=0)$ | 1.0 | 1.0 | 1.0 | 1.0 | 1.0 | 1.0 | 1.0 | 1.0 |
| $C_1(t=0)$ | 1.0 | 1.0 | 1.0 | 1.0 | 1.0 | 1.0 | 1.0 | 1.0 |
| $C_2(t=0)$ | 1.0 | 1.0 | 1.0 | 1.0 | 1.0 | 1.0 | 1.0 | 1.0 |
| $C_3(t=0)$ | 0 or 2.0 | 0 or 2.0 | 0 or 2.0 | 0 or 2.0 | 0 or 2.0 | 0 or 2.0 | 0 or 2.0 | 0 or 2.0 |
| $C_4(t=0)$ | 0 | 0 | 0 | 0 | 0 | 0 | 0 | 0 |

**Table S17.** Model parameters and initial values for the different interaction types (EC = exploitative competition, IC = interference competition, CF = cross-feeding, CD = cross-detoxification and NS = niche separation). For each parameter, the first subscript indicates the species identity and the second (and third) subscript the compound identity. Lines where all parameters are 0 are omitted (e.g. $r_{1,2} = 0$). For $l_{i,j,k}$, subscript $j$ indicates the growth compound on which species $i$ experiences a lag due to compound $k$. Compound $C_3$ is an environmental inhibitor. We change the concentration of $C_3$ to $2.0$ if the growth medium is inhibitory, which prolongs the lag phase before growth on compound $C_2$. Compound $C_4$ is produced by $S_1$ to feed $S_2$ and so is initially always 0.

DRAFT
